# Dark side of the honeymoon: reconstructing the Asian x European rose breeding history through the lens of genomics

**DOI:** 10.1101/2023.06.22.546162

**Authors:** Thibault Leroy, Elise Albert, Tatiana Thouroude, Sylvie Baudino, Jean-Claude Caissard, Annie Chastellier, Jérôme Chameau, Julien Jeauffre, Thérèse Loubert, Saretta Nindya Paramita, Alix Pernet, Vanessa Soufflet-Freslon, Cristiana Oghina-Pavie, Fabrice Foucher, Laurence Hibrand-Saint Oyant, Jérémy Clotault

## Abstract

- Roses hold significant symbolic value in Western cultural heritage, often serving as a symbol of love and romance. Despite their ancient cultivation, the appreciation for the phenotypic diversity of roses emerged relatively recently, notably during the 19th century. This period is characterized by a remarkable expansion in the number of varieties, from around 100 to over 8,000, representing a golden age for roses.
- To trace the history of rose breeding in Europe and unveil genetic changes during this period, we gathered phenotypic and genetic data from 204 accessions. These included botanical roses and varieties cultivated between 1800 and 1910. Whole-genome sequences from 32 accessions were also included.
- Our analysis revealed a temporal shift in the genetic makeup, transitioning from a historical European to a near-Asian genetic background within a few generations. This shift was accompanied by a notable reduction in genetic diversity, attributed to the backcrossing with the less diverse Asian genepool, plus some genomic signatures of selection.
- We have generated the largest GWAS catalog for rose to date, offering a valuable resource for future breeding initiatives. We emphasize the critical importance of preserving ancient rose collections to safeguard diversity and ensure a sustainable breeding for the long term.

## Introduction

The neolithic revolution, the move from nomadic hunter-gatherers to rooted agrarian societies, has profoundly changed the history of humankind (Diamond, 2002). This transition has allowed some populations to have larger food surpluses, favoring rapid population growth, a division of labor, technological innovations and, at the end, the establishment of dominant societies and colonizers (Diamond, 2005). Domestication is expected to generate drastic reductions in the effective population sizes due to the subsampling of the wild progenitor species and the selection pressures that have then further reduced population sizes, which have long been assumed to lead to sudden losses of diversity (Nei *et al.,* 1975; see Gaut *et al.,* 2015 for empirical evidences). Recent investigations thanks to ancient DNA samples, in sorghum in particular, are however more consistent with a long-term gradual, linear or nearly linear, decline rather than a rapid drop, requestioning in part this bottleneck scenario (Allaby *et al.,* 2019; Brown, 2019; Smith *et al*., 2019).

Unlike species domesticated for food production, the domestication of floral plants for aesthetic purposes is assumed to be far more recent, which can therefore even more question the existence of reductions of genetic diversity over these short periods of time. For most of the ornamentals, domestication indeed occurred in the last 500 years (Purugganan, 2022). Economically secure groups, especially the bourgeoisie in search of luxuries or aesthetic pleasure, have played a major role in this advent, leading to an exploding number of domesticated ornamental species, in such a way that ornamentals have been described as outnumbering all other domesticated species combined today (Gessert, 1993, 2010; Chowdhuri & Deka, 2019). A typical example of this new passion for flowers in Western countries is tulips. Initially appreciated by Ottoman sultans and elites, the tulips imported to the Netherlands have led to a tulipmania, an irrational frenzy during the 17th century. Garden roses represent another emblematic example of domesticated flowers (Altman *et al.,* 2022), with a unique symbolic charge (Goody, 1993). Although rose domestication can be traced back as 5000 years BP (Altman *et al*., 2022), the process of domestication is poorly understood. Its golden age is assumed to be even more recent than tulips, dating back to the 19^th^ century (Leus *et al.,* 2018).

Despite roses being cultivated since antiquity, both independently in China and the Mediterranean region, the number of varieties has remained particularly limited for a long time. In ancient Rome, six cultivated species, assumed to correspond to cultivated forms of *Rosa alba*, *R. gallica*, *R. damascena*, and *R. moschata* were described (Pline l’Ancien, 2013). During the Middle Ages, roses were cultivated not only for their ornamental uses but also as religious symbols and for their medicinal properties (Touw, 1982). In Asia, roses were cultivated as early as the Han Dynasty, over 2,000 years ago, with selection of roses with new traits such as continuous flowering. The introduction of Asian roses to Europe at the end of the 18th century revolutionized rose breeding bringing traits such as recurrent blooming, novel scents and new colors (Wylie, 1954).

Thanks to considerable literature review efforts focusing on the 19th century, the history of a spectacular increase - by a factor 80 or so - in the number of rose varieties has been revealed (∼100 in 1800, 6,000-8,000 in 1900, 30,000-35,000 today; Marriott, 2003; Oghină-Pavie, 2021). The diversification during the 19th century is associated with crosses between the two previously isolated genetic backgrounds, the Asian and European gene pools, giving rise to a large number of horticultural groups, including hybrid tea roses from the late 19th century (Martin *et al.,* 2001). Hybrid tea roses are considered as the parents of the modern roses and are known to be phenotypically diverse, harboring traits that originate from Chinese roses, especially the capacity of recurrent flowering, a particularly targeted trait that led to an extension of the flowering period in Europe (Marriott, 2003; Soufflet-Freslon *et al.,* 2021). Large-scale genetic investigation based on more than a thousand varieties genotyped at 32 SSR markers have indeed identified 16 genetic groups, most associated with a gradient of population structure from a European (groups 1-3) to an Asian (group 9) genetic background during the 19th century, with more recent modern roses (>1914) exhibiting a close to Asian genetic structure (group 8, Liorzou *et al.,* 2016). Despite the progress made regarding the description of the genetic structure, there are many unknowns, including the number of generations of breeding, the levels of diversity and their evolution or the proportion of the respective European and Asian genomes. Similarly, while progress has been made regarding the description of genetic variation at some key genes, including *RoKSN* or *AP2*, controlling the duration of the blooming period and the doubling of the petal number, respectively (Iwata *et al.,* 2012; François *et al.,* 2018; Gattolin *et al.,* 2018; Soufflet-Freslon *et al.,* 2021), the genetic bases of most targeted traits in roses remain unknown.

Roses represent an excellent ornamental model to reconstruct the past history of breeding and investigate the genomic footprints of artificial selection. First, and most importantly, roses are maintained and reproduced through vegetative culture, in particular grafting, allowing the continuous maintenance of ancient varieties. This allows the phenotyping of roses that were bred at different periods of time in a single environment. In addition, it allows easy access to fresh material and therefore modern DNA from these varieties, without requiring expensive and challenging strategies such as the use of ancient DNA samples. Second, several reference genome assemblies are available for roses, in particular from the variety *Rosa chinensis* ‘Old Blush’ (Hibrand-Saint Oyant *et al.,* 2018; Raymond *et al.,* 2018), one of the four main ancient Asian roses used in plant breeding history (Liorzou *et al.,* 2016). Third, roses have relatively small genomes, around 500-550 Mbp - which for instance contrasts with the 34-Gbp of the tulip genome - allowing the resequencing of some rose accessions at affordable cost. Fourth, other molecular resources are available for roses, including a high density SNP array (68,893 markers, Koning-Boucoiran *et al.,* 2015), making high resolution mapping and Genome-Wide Association Studies (GWAS) possible (SchulzDietmar *et al.,* 2016; Hibrand-Saint Oyant *et al.,* 2018). However, one challenge associated with the model is the variable ploidy level among roses, ranging from diploid to decaploid, even if most roses are either diploid or tetraploid. GWAS methods were initially developed for diploid models, but more recent developments, especially for polyploids (*e.g.* Rosyara *et al.,* 2016), provide new opportunities to uncover the genetic bases for important traits in roses. One additional difficulty associated with this model is the complexity of the genetic relationships in the *Rosa* genus, with a hundred or more wild species (Wissemann & Ritz, 2007; Debray *et al.,* 2021), for which approximately fifteen of them may have genetically contributed to present-day rose diversity.

In this study, we reconstruct the history of the rose garden breeding in Europe. To do so, we assembled a unique dataset, composed of large phenotypic data, genetic data from 204 roses genotyped with a 69k SNP array, plus whole-genome sequence data from 32 roses. The sequenced accessions are either botanical roses or varieties bred in Europe between 1800 and 1910, and include star varieties at that time. Botanical roses are considered as wild species, but they are maintained in rose collections and are assumed to be non-intentionally selected. By combining population genomics and quantitative genetic analyses, we (i) described the genomic makeup of the rose varieties, (ii) provided an approximate number of generations of selection, (iii) estimated the levels of genetic diversity and investigated their evolution throughout the period, (iv) identified potential footprints of artificial selection, and (v) performed the largest GWAS to date to uncover the genetic bases of many horticulturally important traits.

## Materials and methods

### Sampling & DNA extraction

Roses suspected to be diploid and tetraploid, based on prior knowledge (Liorzou *et al*., 2016), were prioritized during our sampling campaigns (see below). Our sampling also had the objective to encompass a range of rose groups covering the 19th century, including the one that are hereafter referred to as ancient Asian and European roses, early European × Asian hybrids, hybrid tea and botanical roses.

A total of 288 samples were collected in the ‘Rose Loubert’ rose collection and nursery (Roseraie Loubert, Gennes-Val-de-Loire, Pays de-la Loire, France) in summers 2020 and 2021, except for *Rosa* × *wichurana* and ‘Old Blush’, both collected in an experimental collection at INRAE (UE Horticulture, Angers, Pays de la Loire, France).

In complement, we performed whole-genome sequencing for fifteen roses. Newly sequenced samples were chosen, based on their presumed importance during the 19th century (considered as “star varieties” based on extensive literature searches on historical catalogs of varieties; Cristiana Oghina-Pavie, personal communication) and the within-group diversity previously observed at 32 SSR markers (Liorzou et al., 2016). Fresh plant material was collected in the Roseraie Loubert in summer 2021, with the notable exception of ‘Jin Ou Fan Lu’, a cultivar that was previously collected by Shubin Li in China (Liorzou *et al*., 2016). It should be noted that we had access to only a small part of the ancient roses, both because of the investigated collection, but also because a part of the ancient varieties have not been preserved down to this day. As a consequence, our sampling was potentially impacted by a conservation bias.

For each rose, 100 mg of leaves were collected and DNA was extracted from either lyophilised or fresh leaves using the Qiagen 96 Plant Kit (Qiagen, Hilden, Germany). DNA quality and quantity were checked with the Nanodrop technologies and Qubit kit prior to sending samples to the different facilities.

### Genotyping and sequencing data

Genotyping was performed on 3*96-plates at the Gentyane facility (INRAE Clermont-Ferrand, France) using the Affymetrix Axiom technology. To further check the quality of the genotyping (see below), we included technical replicates (21 duplicate samples, as well as a triplicate one, shown in yellow in Table S1).

Regarding sequenced samples, library preparations and sequencing were performed at the Vienna BioCenter Core Facilities (VBCF, Vienna, Austria). For the library preparations, the facility used the Ultra II DNA kit for Illumina (New England Biolabs), which included an enzymatic shearing as a first step. Quality control was performed prior to sequencing. Quality control of the obtained NGS libraries was then assessed prior to the sequencing using the Fragment Analyzer technology. The 15 genotypes were then sequenced on a single NovaSeq SP flow cell (300 cycles, 2*150 reads). Demultiplexing was also performed by the VBCF facility.

### Phenotyping

All phenotyping data were collected in the rose garden (Roseraie Loubert, Rosiers sur Loire, Gennes-Val-de-Loire, France). The number of petals was scored on at least two and at most five flowers. We used averaged petal counts across the different flowers. We also recorded the number of flowers per inflorescence and categorized roses by the primary color of their flowers. Additionally, we measured the number of prickles and acicles, along with several architectural traits: height (meters), circumference (meters), growth form, and bushiness (binary). While pruning is performed within the rose garden, it is important to note that it respects the natural form of roses and is not expected to considerably alter their architecture or growth patterns. However, this limitation associated with the size of the roses in the rose garden should be kept in mind when interpreting these results.

Plants were scored in the rose garden every week during the blooming period (from spring to autumn) for the recurrent blooming. This flowering monitoring was conducted over a period of three years (2014-2016). We have previously developed a statistical method on the flowering curve using Gaussian mixture models and created indicators, as Reb.Mag for the magnitude of reblooming (for details, see Proïa *et al*., 2016). A value of 0 means that the rose flowers only in spring (once-flowering roses), the higher the value (above 0), the higher the reblooming capacity (see Supplementary Note 1). For the sake of clarity, the index shown in Fig. 1B corresponds to a simplified version, with values corresponding to three phenotypic classes (1: non-recurrent flowering, 2: partly recurrent, 3: recurrent flowering), an average value of the index across the three years of scoring was then computed for each individual.

**Figure 1:**
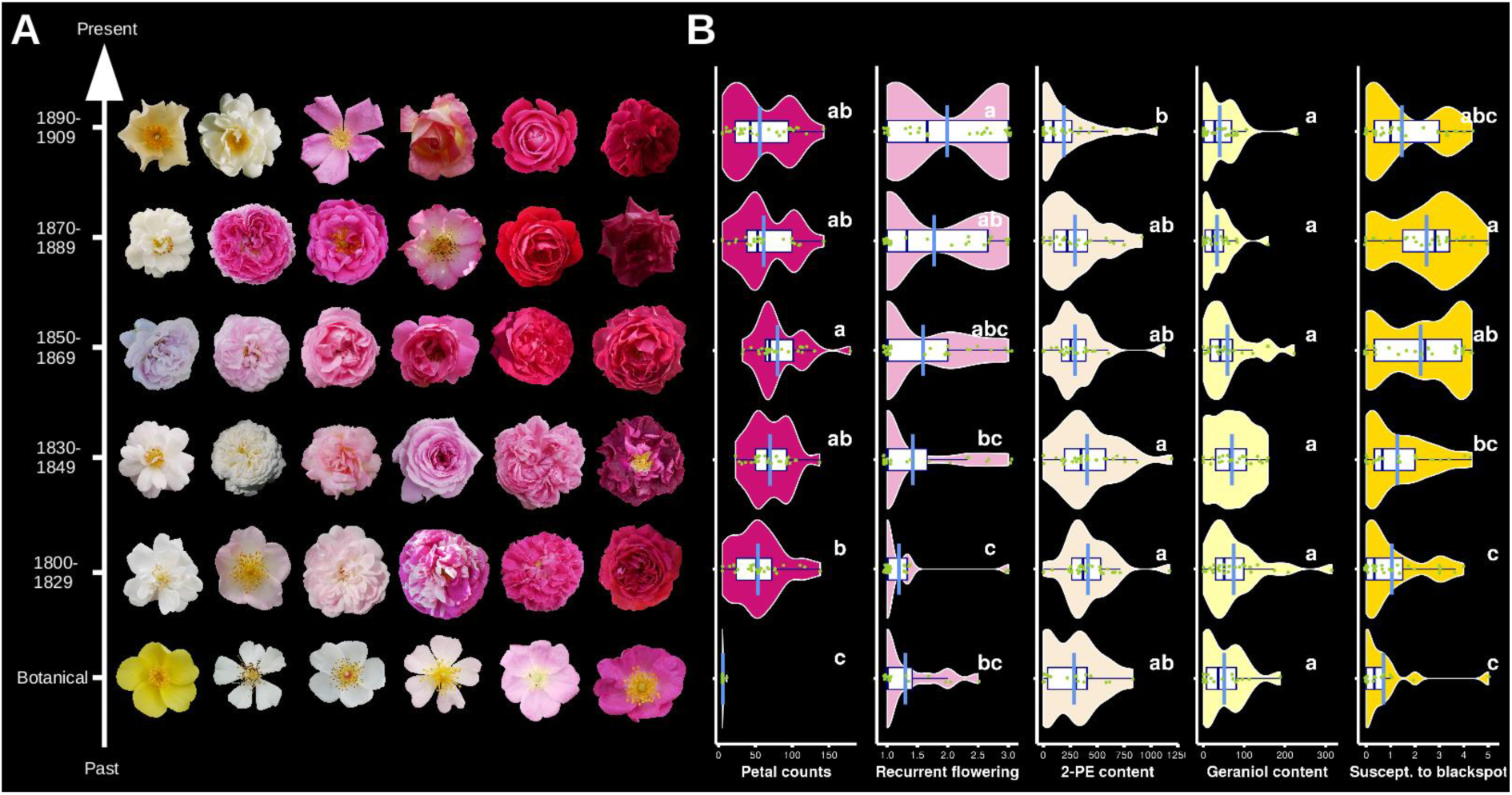
Main phenotypic changes along the 19th century plant breeding history. **A.** Pictures highlighting the phenotypic variation of flowers at different periods of the rose breeding history among the different accessions used in this study. Pictures were taken by T. Leroy in the Loubert rose collection in 2021 and 2022 and selected to be representative regarding the number of petals and the variation of colors observed among all the varieties of a given period. Pictures are not to scale. **B.** Phenotypic variation for some of the studied traits: number of petals (dark pink), recurrent flowering (light pink, estimated by an index), scent components (2-phenylethanol and geraniol contents, light beige and light yellow, respectively) and susceptibility to black spot disease (yellow). Averages are shown with light blue lines. Tukey’s honestly significant difference (HSD) tests at ⍺=0.05 are indicated with letters.

Flowers were also collected during two years (2017 and/or 2018) to analyze scent components. Concretely, from two to six flowers per sample and year (2017: mean=2.34, sd=0.78; 2018: mean=2.33, sd=0.81) were harvested around 10:00 am at stage 4 of flower development, as previously described (Bergougnoux *et al.,* 2007) and volatile compounds were extracted following Roccia *et al*. (2019). Per-genotype averaged values of each compound per year were used for all the analyses.

Susceptibility to black spot disease was also scored in the field over three years (2014-2016). Each cultivar was scored, between July and September, depending on the development of disease and according to the scale indicated in Marolleau *et al*. (2020). The observed black spot symptoms were due to natural infection with natural inoculum of fungal strains. It should be noticed that we were scoring roses in the field, with present-day strains. The strains present at the time of breeding (*i.e.* 19th century) might have been different, including their virulence gene repertoires and aggressiveness factors.

### Data analysis

The analyses performed in the manuscript are described below, considering (i) the analyses based on the SNP array and (ii) those on the whole-genome sequencing data. All the scripts used have been made available on a Zenodo repository (10.5281/zenodo.14450242).

### 1/ SNP array

#### Genotyping

We used a two-round genotyping procedure to ensure highly accurate calls. First, we performed a QC analysis under Axiom Analysis Suite based on all the genotypes (288). We used the Dish Quality Control (DQC), the recommended QC metric for the Axiom SNP array to capture the extent to which the distribution of signal values is separated from background values, with 0 indicating no separation and 1 indicating perfect separation. Based on this genotyping, four individuals exhibited very low DQC (<0.25) and were excluded. Based on all the other individuals (with DQC > 0.85, 284 samples), we generated a first genotype matrix with FitPoly (Voorrips *et al*., 2011) assuming a ploidy=4 for all samples.

Given that roses are grafted, all samples with a close genetic proximity with the rootstock cultivar used in the Loubert’s rose garden were considered as sampling errors and were excluded from the dataset (seven in total). In addition to these previously excluded genotypes, we excluded seven additional genotypes with a DQC < 0.94 and a QC Calling rate < 0.89, resulting in a dataset of 270 individuals. For this new dataset, we performed detailed QC and then converted the output in a FitPoly format thanks to FitPolyTools (version 1.1; unpublished R package by Roeland Voorrips). To take into account the variable ploidy level between individuals, with the vast majority of the accessions used expected to be either diploid or tetraploid, we generated a final genotyping matrix using FitPoly assuming ploidy=4 for all individuals, which means that diploid individuals with heterozygous alleles are generally expected to be called AABB. The rose SNP array targets 68,893 SNPs, with probes for both strands, leading to a total number of 137,786 probes (for details, see Koning-Bourcoiran *et al.,* 2015). After SNP filtering, we obtained a dataset of 92,007 probes, corresponding to 56,467 of the 68,893 unique SNPs. Given all the difficulties associated with the genotyping of the SNP array, especially in a species with a variable ploidy level, we made the choice of using all probes satisfying our QC criteria in the subsequent analyses.

#### QC checking and clone exclusion

Given that roses are clonally propagated and used commercially worldwide, several varieties can be registered under different cultivar names. In addition, spontaneous somatic mutants (also called sport) exhibiting different traits (*e.g.* flower color or shape) are frequent in roses and could have therefore been registered with two different names. To exclude potential clones in the dataset that could bias our quantitative and population genomics analyses, we computed the proportion of similar calls between all pairs of individuals and considered all samples exhibiting a value higher than 0.965 as potential clones. To define this value, we considered the observed distribution of the proportion of similar calls based on all pairwise comparisons and the lowest observed value for a replicate of the analysis (0.973 for ‘*Persian yellow*’), see Fig. S1 and Table S1 for a list. As a consequence, our “clone-corrected” dataset contains 204 unique samples. All subsequent analyses are based on this dataset.

#### Population structure

We used the R package adegenet (Jombart, 2008) to explore the population structure among the 204 individuals. We first used plink (v2.00, Purcell *et al.,* 2007, with --index-pairwise 20 5 0.2) to generate a pruned SNP set of 11,848 SNPs among the 92,007 markers. We then used the df2genind (with ploidy=4) to generate the input files before using the dudi.pca function of the adegenet package. PCA results were then plotted using both the ggplot2 and ggrepel packages (Wickham, 2016; Slowikowski, 2023). To confirm the population structure observed with the PCA, we also used faststructure (Raj *et al.,* 2014), with K ranging from two to five groups.

#### GWAS

Before performing GWAS analyses, we further filtered the dataset in order to exclude poorly informative markers. SNPs with a missing rate greater than 10% and/or a Minor Allele Frequency (MAF) < 0.05 were discarded. GWAS was then performed with GWASpoly (Rosyara *et al*., 2016) for all traits. Concretely, we used the Q (population structure) + K (relatedness) linear mixed model, providing our own Q matrix based on the Principal Coordinates Analysis (dudi.pco, adegenet, Jombart, 2008) as covariates. Considering both the tetraploidy and the absence of strong expectations regarding the additive *vs.* dominant effects of the genes for the studied traits, we considered all the potential models of gene action in polyploids and implemented in different models under GWASpoly (*i.e.* general, additive, diplo-additive, diplo-general, simplex and duplex-dominant models, for details regarding the models, see Rosyara *et al.,* 2016). Concretely, by utilizing GWASpoly to analyze the genetic basis of traits in European roses, our objective was to address the complexity of the polyploid gene action as much as possible. To explore the results, we first generated the QQplot and the Manhattan plot using R base, but we then generated linear or circular representations of the results under the R packages qqman (Turner *et al.,* 2018) or circlize (Gu *et al.,* 2014), respectively. In addition to the Manhattan plots, these circular views include density curves in order to identify some potentially interesting regions, even when the empirical background noise is high, generally associated with a general inflation of p-values (as tracked with the QQplots). Given the number of traits analyzed, as well as the number of models explored for each, all the ‘circlize’ visualizations were made available thanks to a dedicated website (https://roseGWASbrowser.github.io/). The website was successfully tested on various web browsers, including Firefox (*e.g.* v. 133.0) and Chromium (*e.g.* v. 131.0.6778.85). The rest of the results including the p- and q-values, as well as the QQplots and the linear Manhattan plots were made available on a Zenodo repository (see the data availability section).

### 2/ Whole-genome sequences

#### Mapping and calling

A total of 32 genotypes are used as part of the whole-genome analyses, corresponding to a set of 17 rose sequences publicly available and 15 new sequences generated in this study. The new sequences have been made available on the Sequence Read Archive (SRA) under the BioProject PRJNA997103 (see Table S2 for all SRA accession numbers).

Briefly, we used Trimmomatic (v.0.38, Bolger *et al.,* 2014) to remove adapters, trim and filter reads using the following set of parameters: LEADING:3 TRAILING:3 SLIDINGWINDOW:4:15 MINLEN:50. All trimmed reads were then mapped against a high-quality rose reference genome (Hibrand-Saint Oyant *et al.,* 2018) with BWA mem2 (v. 2.2, Vasimuddin *et al.,* 2019) using default settings. Most of the mapped data corresponds to paired ends reads but we also mapped single end reads (*i.e.* corresponding to unpaired after trimming). MarkDuplicates (Picard v. 2.20.7; Picard toolkit, 2019) was then used to remove potential PCR duplicates. Variant calling was performed on GATK4 (v.4.2.2.0, Poplin *et al.,* 2018), following the GATK best practices guidelines. In brief, we performed a first round of HaplotypeCaller to generate the GVCFs, followed by a CombineGVCFs and a GenotypeGVCFs with the “--all-sites true” option. Variants were then filtered with VariantFiltration, discarding variants with SOR>4.0, MQ<30, QD<2, FS>60, MQRankSum< -20, and ReadPosRankSum either <- 10 or >10.

#### Ploidy inference

The ploidy level of each sample was empirically determined by considering the observed allelic balance at heterozygous sites. Specifically, for each individual, we subsampled 2% of the total number of SNPs and then considered, for each individual, the heterozygous calls exhibiting a coverage greater or equal to 20 and then generated the per-individual density curves of the allelic balance based on all these SNPs. Because most of the polymorphisms were expected to be rare in the population, we could observe the first neat peak to be near f=0.25, 0.33 and 0.5 for tetraploids, triploids and diploids (see Fig. S2). Similar methods have been developed over the last decade to empirically estimate the ploidy level (*e.g.* ploidyNGS, Augusto Corrêa dos Santos *et al.,* 2017). To ensure that the inferred ploidy level was sufficiently robust for our subsequent analyses, we checked the overall consistency of the results with previous in-house estimates of the ploidy level (for details, see Liorzou *et al.,* 2016 and http://dx.doi.org/10.5281/zenodo.56704).

#### Population structure & kinship

Population genetic structure among the 32 varieties was performed using Principal Component Analysis (PCA). Analyses were run on a set of 17,669 SNPs, which was generated by first randomly selecting 50,369 biallelic SNPs across the genome and then pruning this set for linkage disequilibrium with plink (v1.90, parameters: “--index-pairwise 20 5 0.2”). We used the snpgdsPCA function from the SNPRelate R package (Zheng *et al.,* 2012) to generate the PCA. We then used ggplot2 (Wickham, 2016) to generate the final plots. Given that the reference corresponds to an ancient Asian rose, we also empirically checked that the observed population structure could not have been generated by a strong mapping bias (Fig. S3). Similarly, given that the first round of calling assumed diploid individuals which is not correct for all individuals (see ploidy inference), we also performed PCA directly based on allele counts (AD field from the vcf) from a list of 10.5 million SNPs randomly selected among the biallelic SNPs to evaluate the robustness of our population structure (i) to a larger SNP set and (ii) to deviation from diploidy. PCA was performed with the function ‘randomallele.pca’ from the R package ‘poolfstat’ (Hivert *et al*., 2018; Gautier *et al*., 2022; Supplementary Note 2). In addition, as a control of the population structure observed using PCAs, we also used faststructure (Raj *et al.,* 2014) on the same 50k SNP set. To do so, we first used plink (v2.00, Purcell *et al.,* 2007) to generate the input files and then inferred the population structure for K=1 to K=6. Barplots of the inferred individual ancestry memberships were also generated using ggplot2.

We performed a family relationship inference by estimating kinship coefficients under KING (Manichaikul *et al.,* 2010) based on the whole SNP set (77,862,879 filtered SNPs). The initial objective of this analysis was to ensure that no clones were present in our sequencing dataset (see also above). But given that KING is able to infer relationships up to the third-degree (*i.e.* great grandparents and great grandchildren, first cousins, or similar), we used KING to infer some potential family relationships. We first generated the input files with plink (v2.00) and then estimated kinship coefficients. Kinship values higher than 0.354 correspond to clones, while coefficients coefficients within the intervals [0.177, 0.354], [0.0884, 0.177], and [0.0442, 0.0884] correspond to 1st-degree, 2nd-degree, and 3rd-degree relationships, respectively. Since KING analysis was conducted on the initial SNP calling round, resulting in diploid calls, we cannot rule out a potential impact of variable ploidy levels among the roses in the accuracy of the number of generations (Supplementary Note 2). We used the R package igraph (Csárdi & Nepusz, 2006) to generate a comprehensive network based on all the inferred family relationships.

#### Nucleotide diversity estimates

As compared to population structure and kinship inferences, estimates of diversity are highly dependent on the final quality of the SNP set. To ensure unbiased estimates, we used the filtered set of SNPs based on the 32 samples (77,862,879 filtered SNPs) to perform a Base Quality Score Recalibration (BQSR) on the 19 retained samples, as recommended by GATK in the absence of a highly-reliable SNP catalog for the focal species. Then, we used GATK’s ApplyBQSR function. HaplotypeCaller was then performed assuming the inferred ploidy (see previous section) using option “--ploidy”, before using CombineGVCFs, GenotypeGVCFs and VariantFiltration as previously described to generate a final SNP set of 54,481,222 filtered SNPs. We then reconstructed whole-genome fasta sequences based on the complete VCF (vcf with variant and non-variant positions; to know more about the importance of using non-variant positions, see Korunes & Samuk, 2021) following Leroy and collaborators (2021a, 2021b). The approach was adapted to the rose specificities to reconstruct up to 4 sequences per individual, by explicitly considering the inferred ploidy (per-individual number of alleles in the GT field). Briefly, the pipeline reconstructs the sequence by considering the coverage at each position. All positions that are not in between the 5th and 95th centiles of the individual coverage, as well as those with coverage lower than 3 were hard-masked in the reconstructed sequences. For positions satisfying the coverage criteria, the reference allele is added to the reconstructed sequences, except at PASS positions, where alleles are added according to the genotyping calls (*i.e.* the GT field). Triallelic (or more) variants were discarded. Similarly, indels were ignored to keep the sequence length the same across individuals and therefore obtain perfectly aligned sequence blocks. Nucleotide diversity and Tajima’s D were then computed on non-overlapping 100-kbp sliding windows following Leroy *et al*. (2021a) for each group, based on the individuals falling in the four following groups: ancient Asian, ancient European, early European x Asian and hybrid tea roses (Table S2). Botanical roses were not included in this part of the study because their population structure is not consistent with a single group.

A Reduction of Diversity (RoD) index was computed by 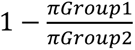, with *Group1* and *Group2* corresponding to hybrid tea roses and ancient European, respectively, for a comparison aiming at estimating the evolution of the diversity in hybrid tea roses as compared to the ancient European samples. A positive value of the RoD index therefore indicates a net reduction of the nucleotide diversity between the two groups. Circular visualization was generated using the R package circlize (Gu *et al.,* 2014).

#### Diagnostic alleles

We then considered allele frequency at diagnostic alleles between the ancient Asian and European genetic backgrounds as a source of information regarding the local ancestry of early European x Asian and hybrid tea roses. For each filtered SNP among the final list of 54,481,222 SNPs, allele frequencies of the four groups were computed directly based on the filtered joint vcf thanks to a home-made script (script_freqgroup4localancestryprop.py), which was made available on a Zenodo repository (DOI: 10.5281/zenodo.14450242). Given that the number of genotypes available per group is limited (4-7 individuals, corresponding to a total of 16 chromosome sets per group), allele frequency was only computed if the number of missing alleles was lower or equal to 4 (*i.e.* corresponding to a maximum of 1 tetraploid individual or 2 diploid individuals without genotyping calls). We then subsampled SNPs exhibiting diagnostic alleles, defined as those for which the reference allele frequency is 0 in one group and 1 in the other. In this specific example, all diagnostic SNPs identified - at the notable exception of 2 SNPs that were subsequently excluded - exhibit an allele frequency of 0 in the European and of 1 in the Asian group. Indeed, all the raw sequencing data were mapped against the genome of an ancient Asian genotype (‘Old Blush’; Hibrand-Saint Oyant *et al.,* 2018). Consequently, the reference alleles are expected to be more associated with the ancient Asian background. The match is almost perfect here given that we also used the ‘Old Blush’ variety as a member of the ancient Asian group in our study. As a consequence, all diagnostic alleles are expected to have an allele frequency of 1 in the ancient Asian group and, consequently, 0 in the ancient European group. Importantly, the diagnostic SNPs are based on our limited panel of varieties. Consequently, all are not expected to be truly diagnostic based on a larger panel.

Local ancestry of early European x Asian or hybrid tea groups was then estimated by computing the median of the observed allele frequencies at diagnostic markers. Estimates were based on non-overlapping 100-kbp sliding windows spanning the whole genome for windows containing at least 5 diagnostic SNPs.

#### Local footprints of selection

To know more about the genomic regions targeted by 19^th^ century breeders, we identified regions that could be consistent with selective sweeps. One limitation associated with the model is the recent history of the rose breeding and the associated limited number of generations of recombination (see results). As a consequence, explicit selective sweep methods could not be used here to identify potential footprints of selection. As an alternative, we identified regions simultaneously exhibiting the lowest π, the most negative Tajima’s D and the highest RoD values. Given that this strategy does not fully circumvent the problem of the lack of generations of recombination, we have decided to focus our detection on hybrid tea roses, the latest group in the history of breeding among those investigated in this study, and therefore expected to have experienced the highest number of generations of recombination, in order to deliver more interpretable results. We considered two sets of candidate windows. The stringent set corresponds to windows falling in the last centile of the three metrics supporting a potential selective sweep (*i.e.* bottom 1% of nucleotide diversity and Tajima’s D values, top 1% of the RoD comparing hybrid tea roses with either Ancient European or Ancient Asian), and a less stringent set considering the last five centiles (Table S4).

## Results

### 19th century: directed or unintended evolution of some key rose traits

Asexual rose propagation through grafting or cutting allows the maintenance of ancient roses in gardens and therefore the direct comparisons of phenotypes obtained at different periods of the breeding history. We explored the phenotypic variation of a large collection of botanical and ancient accessions, representing more than 200 roses bred between 1800 and 1910 (Fig. 1A), a period which is considered as the golden age for rose breeding. In this study, we investigated about 30 traits in 200+ varieties, including several presumably crucial traits for breeding, such as the blooming period, the color and number of petals, the number of prickles, the resistance to black spot disease, plus some flower and plant architecture phenotypes (Fig. 1B). A particularly massive effort was made to analyze the floral scent by quantifying the volatile compounds of the collection using a GC-MS strategy, including the 2-phenylethanol and the geraniol, which represent two main components of the floral rose scent (Spiller *et al.,* 2010; Magnard *et al.,* 2015; Roccia *et al.,* 2019; Caissard *et al.,* 2022).

Among all these traits, those associated with flowering are expected to be the main target of selection during the 19th century, including the production of roses containing tens of petals and the ability to have an extended blooming period. Most botanical accessions indeed produce flowers containing only five petals (Fig. 1B) and during a limited period of time in spring. Based on our sampling, the mean number of petals has strongly increased during the first half of the 19th century (including roses with up to 210 petals), emphasizing the importance of selecting for this trait at the beginning of the 19th century. From 1870, the trend reversed with a reduction of the mean number of petals. Phenotypic variation also supports a relatively constant increase of the repeat-flowering phenotype based on our rose collection, with a progressive increase of the mean of the recurrent blooming index during the 19th century (Fig. 1B).

A main criticism associated with modern roses is the popular impression of a reduction of the flower fragrance. Based on our collection of roses, a slight reduction was indeed observed during the 19th century based on the mean contents of both 2-phenylethanol and geraniol, two main components of the rose scent. Changes during this period are however either weakly significant or non-significant depending on the components, owing mainly to the large observed variance among roses of each period (Fig. 1B).

Temporal changes in the resistance to *Diplocarpon rosae*, the pathogen responsible for black spot disease - one of the most serious diseases on garden roses today -, were also investigated. As compared to the previous examples, resistance to black spot was likely not directly selected by breeders at that time since rose growers started to show interests for *Diplocarpon rosae* (also called *Marssonina* ou *Marsonia rosae* Briosi & Cavara) at the beginning of the 20^th^ century (Ducomet, 1903; Cristiana Oghina-Pavie, personal communication), even though today the resistance to this fungal pathogen represents one of the main breeding targets. Botanical roses from our collection exhibit exceptional resistance levels to *D. rosea* (*i.e.* low susceptibility score) but this resistance has progressively plummeted during the rose breeding history, at least until 1890 (Fig. 1B, see also Fig. S4), which therefore questions the origin of this erosion of resistance (loss of resistance alleles, evolution of the pathogen, etc). Yellow and orange-colored roses are often suspected by breeders to be more susceptible to black spot, hypothetically leading to a trade-off between colors and resistance for breeders. No significant differences in susceptibility were however observed between varieties with different main color petals (Fig. S5). The hypothesis regarding the higher susceptibility of yellow flower varieties remains open since our sampling is biased toward pink and white flowers, largely underrepresenting the genotypes with yellow flowers, mostly because roses with yellow- or orange-colored flowers were relatively rare during the 19th century (Leus *et al.,* 2018).

### A rapid shift from a European to an Asian genetic background

The same large collection of rose varieties was genotyped at 68,893 SNPs using the Rose Genotyping Array (WagRhSNP Axiom Array, Koning-Boucoiran *et al.,* 2015), a SNP array developed from modern cut and garden roses. After the removal of clonal genotypes, a final set of 204 rose varieties was used for the subsequent analyses (see Materials and methods, Fig. S1 and Table S1). The first axis of a Principal Component Analysis (PCA) explains more than a fifth of the total variance (21.4%) and isolates the two ancient genetic backgrounds: an Asian one (in yellow) and an European one (in blue, Fig. 2A). The whole dataset forms a gradient of population structure between the two backgrounds, which is consistent with the frequent crosses between European and Asian roses throughout the 19th century. The second axis isolates some samples, including most of the botanical samples, from the Asian-European gradient. This population structure is mostly consistent with a previous report based on the genotyping at 32 microsatellite markers of an even larger collection of varieties and the identification of 16 genetic groups (Fig. 2A, Liorzou *et al.,* 2016). Independently from this first dataset, we resequenced 15 varieties and collected publicly available rose genomes to generate a dataset of 32 samples, composed of ancient Asian, ancient European, early European x Asian, hybrid tea and botanical samples (Table S2). After joint-genotyping and SNP filtering, we generated a dataset of 77,862,879 filtered SNPs on the 32 samples. PCA based on a LD-pruned set of 17,669 variants among 50,369 randomly selected SNPs (Fig. 2B) is consistent with an Asian-European gradient of population structure, as previously identified with the SNP array (Fig. 2A). The proportion of variance explained by the first axes is however lower based on the sequencing data, likely due to SNP ascertainment bias associated with the design of the array, since the SNP set was established based on the analysis of more modern varieties, therefore emphasizing more on the European-Asian gradient (*i.e.* the axis 1). Based on the WGS data, some axes can be built due to a single botanical individual (*e.g.* axes 2 and 4, Fig. S8), suggesting that a large proportion of the diversity is indeed only present in the botanical accessions.

**Figure 2:**
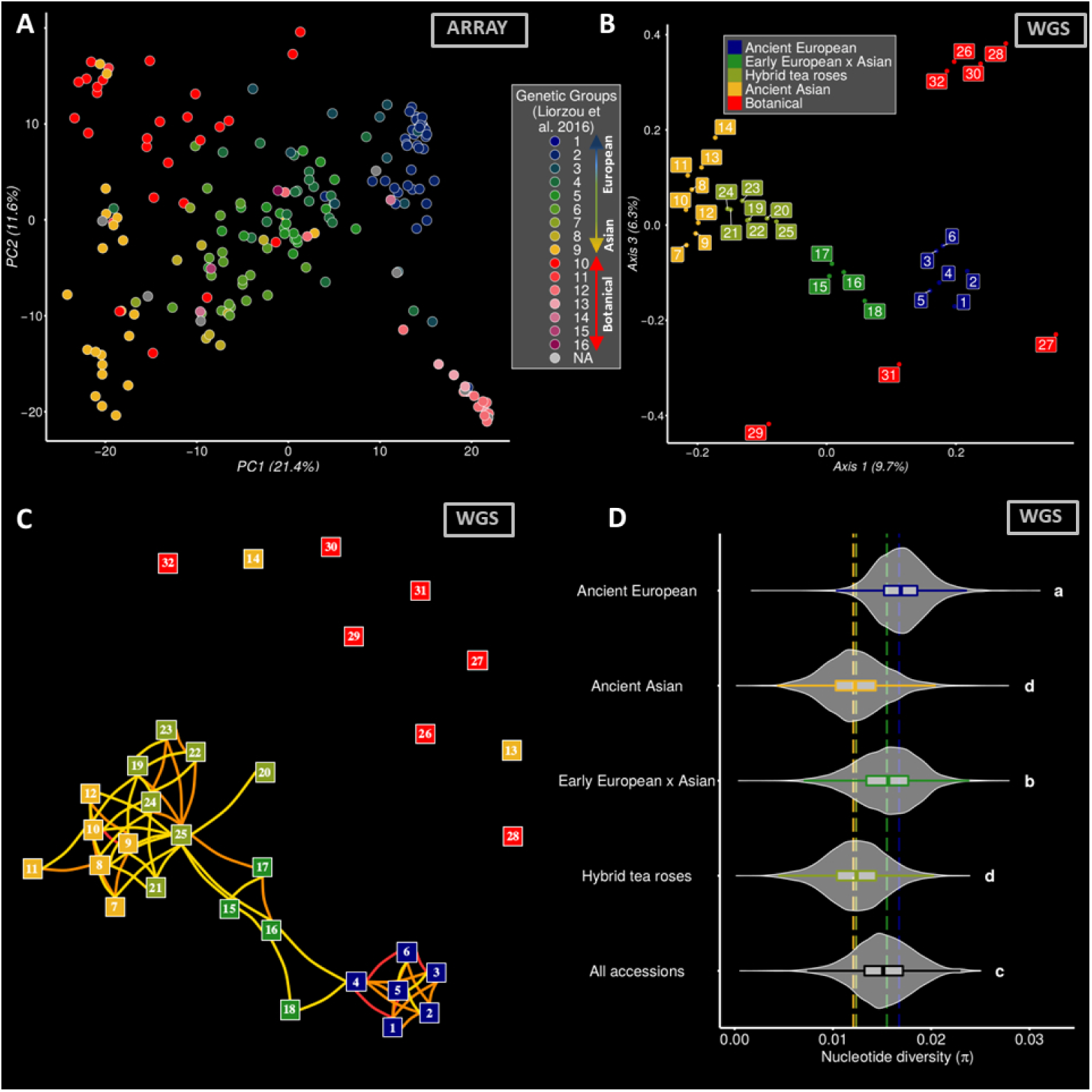
Population structure and levels of genetic diversity in several rose groups used in the 19^th^ century rose breeding history. **A.** Population structure of the 204 varieties genotyped with the rose SNP array based on the two first axes of a Principal Component Analysis (PCA). To check the consistency with previous works, the membership of the samples to one of the 16 genetic groups defined by Liorzou and collaborators (2016) based on the analysis of SSR markers are shown (for details, see Liorzou et al., 2016). Roses with ancient European and ancient Asian backgrounds are shown in blue and yellow, respectively (typically corresponding to groups 1-3 and 9, respectively; see Figs. S6-S7). **B.** Population structure based on the whole-genome sequences used in this study. Axes 1 and 3 of a Principal Component Analysis (PCA) based on a random sampling of 50k SNPs. Ancient European, ancient Asian, early European x Asian, hybrid tea varieties and botanical accessions are shown in blue, yellow, green, khaki and red, respectively. Results from other components as well as more detailed information about the cultivar IDs are shown in Figs. S8 and Table S2, respectively (but see also Fig. S9 and Supplementary Note 2). **C.** Network of the relationships inferred based on the kinship coefficients estimated under KING based on the whole SNP set of 77,862,879 SNPs. First-, second- and third-degree relationships are represented with red, orange and yellow lines, respectively. See Table S2 for a correspondence table between accession numbers and cultivar IDs. **D.** Genetic diversity estimates based on the different groups. Distribution of the nucleotide diversity levels based on non-overlapping 100 kb sliding windows spanning the genome (each dot corresponds to a window). Genomic windows on chr00 are not included. For ease of comparison, a vertical line is shown representing the mean π value per group.

Family relationships were then inferred using KING (Manichaikul *et al.,* 2010) based on the pairwise kinship values calculated on the sequencing data. Among the 32 samples, only a few individuals are not connected to at least another individual of the dataset through a third degree relationship (*i.e.* kinship coefficient > 0.0442) (Fig. 2C). These individuals correspond to all botanical samples, plus two ancient Asian samples (*R. odorata gigantea* and *R. chinensis spontanea,* numbers 13 and 14 in Fig. 2C, respectively). The rest of the individuals are connected through a large network, linking the ancient European and the ancient Asian backgrounds through the early European x Asian and hybrid Tea varieties (Fig. 2C). This result is remarkably consistent with the expected history of selection during the 19th century, with (1) the intercrossing of the Asian genepool with the European one in the European breeding practices, (2) leading to the obtention of early European x Asian varieties, and (3) the backcrossing of these early hybrids with the ancient Asian genepool to obtain the hybrid tea varieties. Obtaining such a network with a so limited number of samples, and without integrating prior knowledge about the potential pedigrees, suggests that the selection was based on a quite narrow diversity, in terms of the number of varieties used. In addition, it suggests that the number of generations of selection was also very limited (but see also some limits in Supplementary Note 2). Hybrid roses from the 19th century are indeed connected to Asian and European gene pools by less than 10 generations of breeding (Fig. 2C, but see also Supplementary Note 2).

To investigate the contribution of Asian or European gene pools to early European x Asian and hybrid tea roses, we identified SNPs with diagnostic alleles between the ancient Asian and ancient European samples. In total, 170,637 diagnostic SNPs were identified among the final list of 54,481,222 SNPs (0.31%), distributed on all chromosomes (Table S3). At these markers, the early European x Asian samples remarkably have a 1:1 ratio, *i.e.* an almost equal share between the two gene pools (50.9% and 49.1% of Asian and European alleles, respectively; Fig. S10). One might propose that this 1:1 ratio implies that all early European x Asian samples used in the study are first-generation (F1) hybrids between the ancient Asian and European groups of samples. However, detailed analyses at the individual level are inconsistent with this hypothesis (Supplementary Note 3). If we can exclude the F1 hypothesis, we can then subsequently exclude the BC1 hypothesis, despite the similarly noticeable 3:1 ratio observed for hybrid tea roses (74.8% and 25.2% of Asian and European alleles, respectively; Fig. S10). To gain a more accurate understanding of the 19th-century rose breeding history, especially concerning the number of generations leading to the “star varieties” in our study, additional research reconstructing pedigrees based on a larger number of whole-genome sequenced accessions will be needed.

### Genetic diversity erosion

Nucleotide diversity was estimated over non-overlapping 100-kbp sliding windows on the four groups (namely, ancient European, ancient Asian, early European x Asian and hybrid tea roses), each containing the same number of complete chromosome sets (16) after considering the inferred ploidy based on the distribution of allelic balance (Table S2, *e.g.* in Ancient Asian the 16 complete chromosome sets corresponds to 6 individuals inferred as diploid individuals, plus a tetraploid). Among all the dataset, the genetic diversity in roses was relatively high with an average π of 1.47×10^-2^ (Fig. 2D). Significant differences in diversity were observed between groups, with higher diversity in the ancient European (1.67×10^-2^) than in the ancient Asian samples (1.20×10^-2^).

The “early European x Asian” group exhibits an averaged nucleotide diversity of 1.52×10^-2^, a value that slightly exceeds the expectation assuming both the previous estimates and the genetic makeup of this group (50.1%/49.1%, expectation: 1.43×10^-2^). This slight burst of genetic diversity in the first generations of hybridization could be explained by the mixing of parental alleles from the two quite divergent genetic backgrounds.

The hybrid tea group has a low nucleotide diversity (average π_observed_=1.21×10^-2^; Fig. 2D), a level that is roughly similar to those of the Asian background (π_observed_=1.20×10^-2^). This diversity is much lower than expected assuming the inferred genomic composition (74.8% Asian, 25.2% European, π_expected_=1.32×10^-2^), corresponding to an average Reduction of Diversity index (RoD) of 8.1%. Given that this breeding history occurred in Europe and that hybrid tea roses have then been preferred to the early European x Asian hybrids in the gardens of residential homes, the evolution between hybrid tea roses and the other European groups is informative about the genetic loss during this period. Comparing these two groups, hybrid teas exhibit an important reduction of diversity as compared to the early European x Asian roses (average RoD=18.5%) and even more as compared to the ancient European accessions (average RoD=27.5%). One could argue that hybrid teas were exported worldwide, including in Asia and therefore that this history of selection could have contributed to an increase of the genetic diversity in Asia. Considering this hypothesis, we compared ancient Asian and hybrid teas and observed such a potential - albeit marginal - gain (RoD= -1.04%). Our results are therefore overall consistent with a net and rapid erosion of nucleotide diversity in Europe during the rose breeding of the 19^th^ century.

The pattern of reduction of diversity between groups is consistent across the chromosomes, even if it varies in intensity (Fig. S11). Comparing the diversity of hybrid teas as compared to ancient European roses, 2.3% of the windows exhibit a reduction of at least half of the diversity (105 among 4,628 windows with RoD exceeding 50%), a value that appears particularly strong given the extremely limited number of generations (Fig. 2C). The maximum observed RoD value on a window is 87.3%, but corresponds to an unanchored scaffold (Chr00) and should therefore be interpreted cautiously. The maximum value for a window anchored on a chromosome is 77.4% (Chr6: 11.2-11.3 Mb). Among the 60 windows with a ROD>0.5 and that are not located on unanchored scaffolds, 35% (21) and 43% (26) are located on chromosomes 3 and 5, respectively, suggesting that these two chromosomes were particularly targeted by artificial selection.

### Detecting local genomic footprints of artificial selection

To know more about the history of rose breeding, we investigated how the genomic landscapes of diversity and Tajima’s D have evolved during the rose breeding history (Fig. 3). All landscapes of nucleotide diversity are highly correlated between groups (Fig. S12), even those of the ancient European and ancient Asian groups (Pearson’s r=0.301, p=3.57e^-87^). However, the Tajima’s D landscapes of these two groups are independent (Pearson’s r=-0.02, p=0.20), suggesting that the correlations of the genomic landscape of diversity are not associated with similar selection targets, but rather more likely shaped by some long-term shared genomic features (*e.g.* the recombination landscape, Shang *et al.,* 2023). The genomic landscapes of diversity become more and more correlated with time, especially when compared to the ancient Asian background (ancient Asian vs. early European x Asian samples, Pearson’s r=0.408, p=1.44e^-165^; ancient Asian vs. hybrid tea samples, Pearson’s, r=0.566, p=0). Consistently, the Tajima’s D landscapes of the hybrid tea is more correlated with the one of the ancient Asian (Pearson’s, r=0.339, p9.26e^-112^) than with the one of ancient European samples (Pearson’s, r=0.170, p=2.70e^-28^). However the high level and significance of the latter correlation also suggest that artificial selection targeted alleles from European origin in some genomic regions.

**Fig. 3:**
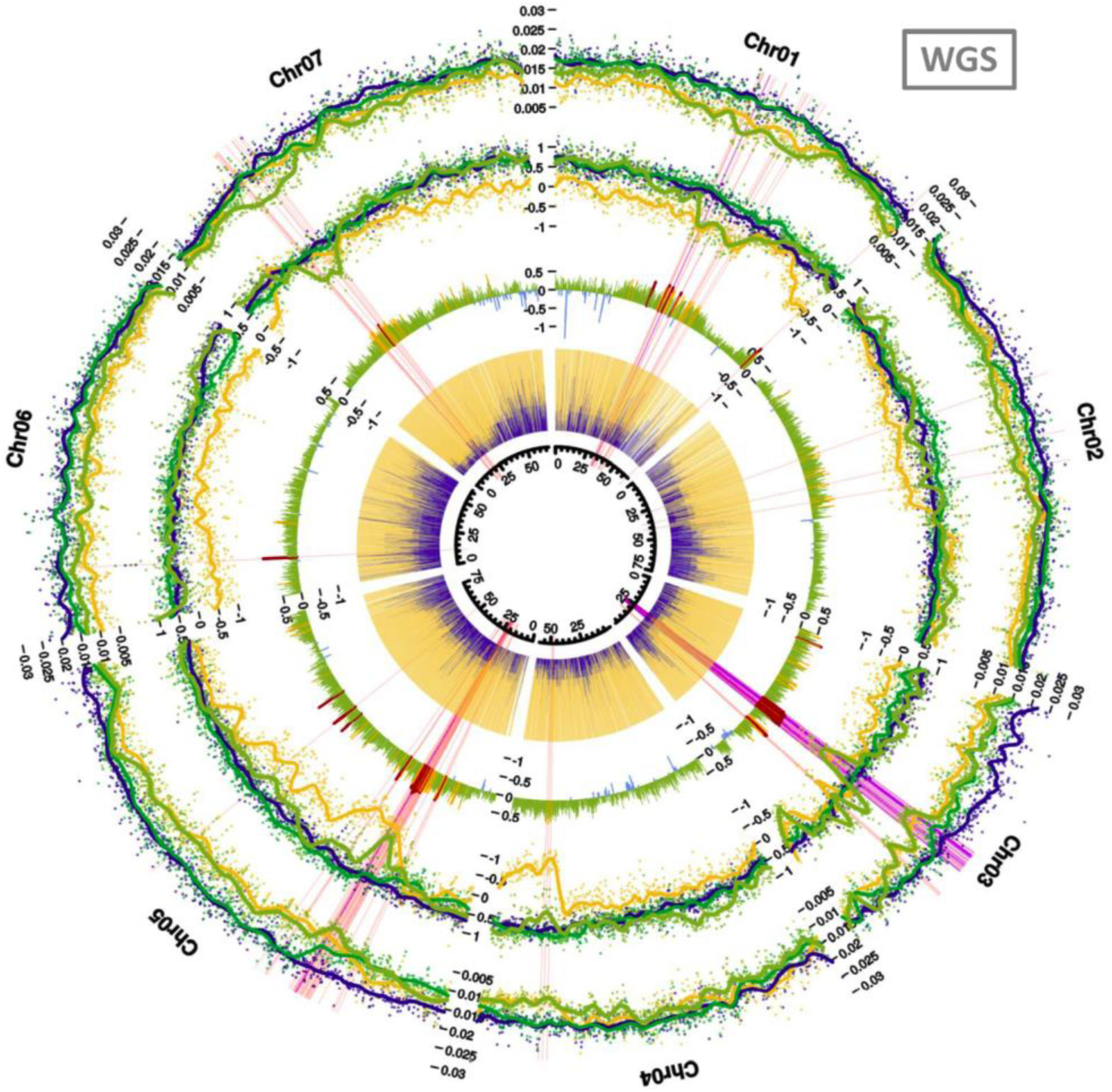
Detection of regions exhibiting footprints of artificial selection in hybrid tea roses. From external to internal: nucleotide diversity, Tajima’s D, RoD _Hybrid tea vs. Ancient European_, and local ancestry as estimated at diagnostic SNPs. Blue and yellow refer to ancient European and ancient Asian backgrounds respectively, while dark and light green refer to early European x Asian and hybrid tea groups, respectively. Pink and purple lines correspond to windows that simultaneously exhibit one of the lowest nucleotide diversity (bottom 5% or bottom 1%, respectively), one of the lowest Tajima’s D (bottom 5% or bottom 1%, respectively) and one of the highest RoD (either in RoD _European vs. Hybrid tea_ or RoD_Asian vs Hybrid tea_, top 5% or top 1%, respectively). Results from chr00 (corresponding to all unanchored scaffolds in Hibrand-Saint Oyant et al., 2018) are not shown.

Interestingly, the genomic landscapes of nucleotide diversity and Tajima’s D are significantly correlated in ancient European (Pearson’s r=0.061, p=9.78e^-5^), as well as in ancient Asian samples (Pearson’s r=0.430, p=2.79e^-185^). However this correlation largely differs in intensity between the two groups. This result is consistent with the lower diversity (Fig. 2D) and the larger variance in Tajima’s D in the ancient Asian group (Fig. S13). All together, these results suggest that ancient Asian samples were more impacted by artificial selection prior to their use in the European plant breeding programs than the ancient European samples. Our genomic evidence could be interpreted with regards to the documented knowledge of the long-term history of rose cultivation in China. Chinese roses have been widely cultivated since the Han Dynasty, and selection is known to have occurred during this period, at least for some important traits, including the extension of the flowering period.

In order to identify some local genomic footprints of selection, we used hybrid tea roses as references. Indeed, given the limited number of generations of breeding detected (Fig. 2C), the use of hybrid teas allows us to maximize the probability of identifying local footprints of artificial selection. Hybrid tea genomes were scanned for windows exhibiting a shared signal of (i) low π, (ii) negative Tajima’s D and (iii) high RoD π_Hybrid Tea_ *vs.* either π_Asian_ or π_European_ (Figs. 3; see Materials & Methods for details). The three summary statistics do not fully overlap in their detection capabilities, each offering distinct specificities for detection (Fig S14). Eighteen windows were detected using the most stringent criteria (last centile of the three metrics, purple lines in Fig. 3), including 15 on a 4-Mb region of the chromosome 3 (from 24.1 and 28.2 Mb, see Table S4). This region is in the immediate vicinity of a previously characterized 5Mb-inversion containing the *RoKSN* gene (28.8 - 33.1 Mb, Kawamura *et al.,* 2022), a presumably main breeding target to obtain recurrent-blooming roses with an extended period of blooming. In addition, to these 18 windows, we detected 81 additional windows (last five centiles of the three metrics, pink lines in Fig. 3). Remarkably, five of these additional windows are located at the very end of the inversion (32.5 - 33.1Mb, see Table S4). The three additional windows detected using the most stringent filtering criteria are located on chromosome 5 (two adjacent windows, between 22.9 and 33.1 Mb) and on chromosome 1 (33.6-33.7 Mb, Table S4). Most of the additional regions based on the less stringent criteria (last five centiles, see Table S4) are consistent with these three previously detected genomic regions (*i.e.* on chromosomes 1, 3 and 5), but also highlighted a fourth region on chromosome 7 (11 windows between 15.9 and 21.1 Mb, Fig. 3 and Table S4).

### The largest rose GWAS catalog to date

In addition to reconstructing the history of rose breeding, we conducted a large Genome-Wide Association Study (GWAS) using GWASpoly (Rosyara *et al.,* 2016) with the aim of identifying associations between the genotypes at the 204 varieties and the different phenotypes, accounting for the levels of population structure and kinship among individuals. All GWAS was performed to identify associations at 46,691 markers, which corresponds to the fraction of markers from the SNP array with a missing rate lower than 10% and a MAF exceeding 0.05. Given the large number of phenotypes we investigated, as well as the variety of scenarios possible regarding dominance in polyploids (Rosyara *et al.,* 2016), a dedicated website portal has been created in order to explore all the results (hereafter referred to as the rose GWAS browser, https://roseGWASbrowser.Z.io/). The online browser allows to select a category of traits (*e.g.* petal counts or colors, plant architecture, scent), then a specific trait (*e.g.* 2-phenylethanol content), and select a specific scenario regarding the polyploid gene action, among additive, simplex dominant and duplex dominant scenarios. By default, the rose GWAS browser reports the most general type of genetic model (“general model”), a model that has the advantages of making no assumptions regarding the underlying gene action, since the effect for each genotype class can be arbitrary (*e.g.* associations due to ABBB genotypes as compared to all others). General models have the advantage of providing an overview of the results for the other models, explaining also their use for Fig. 4. However, as previously indicated, results from the other models, which are more explicit regarding the gene action should be investigated in complement. Given that the number of SNPs also vary along the genome, we also use a local enrichment in associated SNPs to more easily identify associated regions. All the results from the GWAS were also made available on the Zenodo repository, including QQplots, linear Manhattan plots, as well as all p-values and q-values. For some traits, the GWAS explorer not only reports the results of the GWAS when the trait was analyzed quantitatively, but also when analyzed qualitatively, which can be meaningful when distribution of the trait values was observed to be more bimodally distributed.

**Figure 4:**
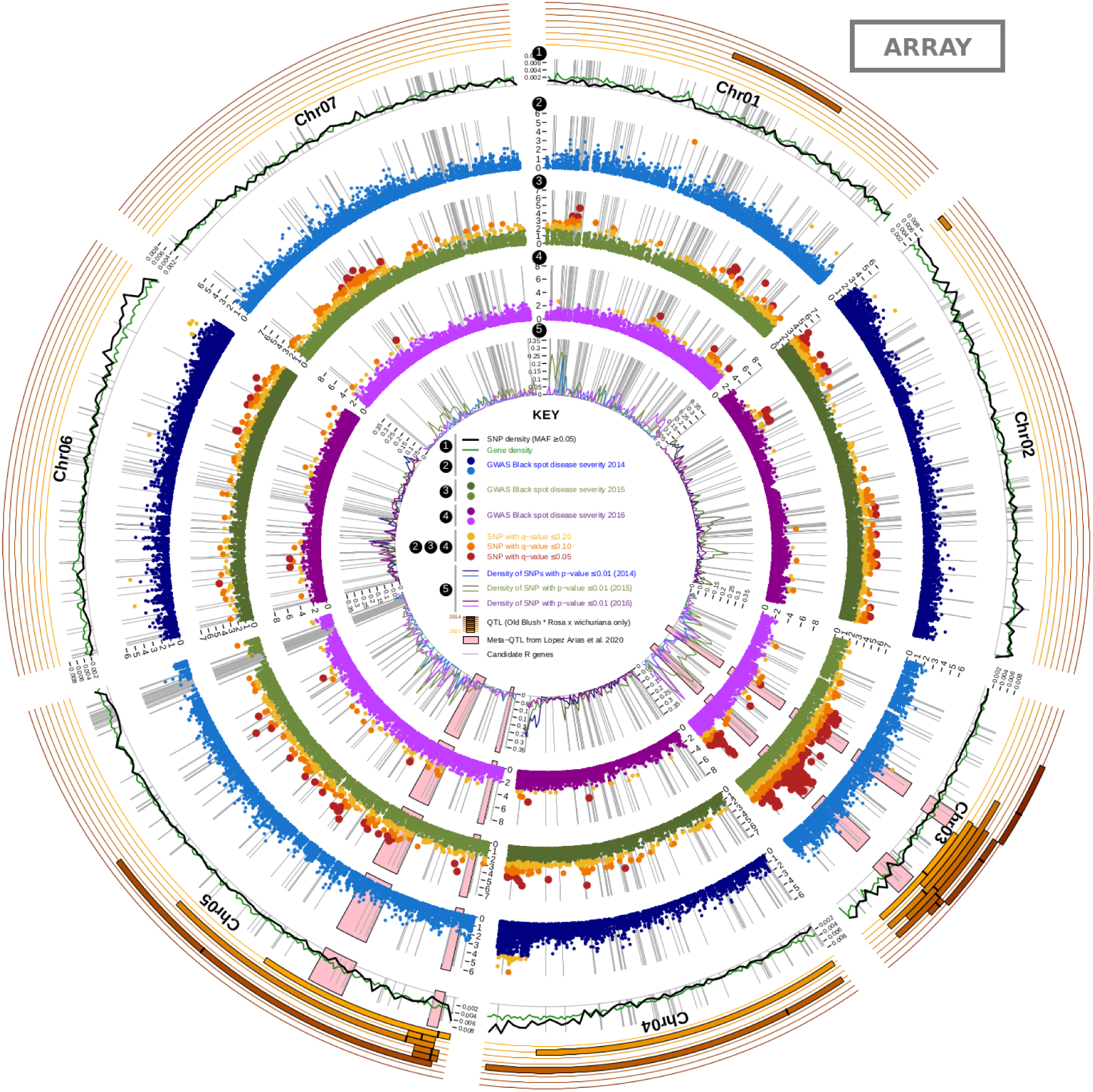
GWAS for black spot resistance. 1: Per-window SNP array density and gene density based on the reference genome (Hibrand-Saint Oyant et al., 2018). 2-4: Manhattan plots of the p-values along the genome for the three years of scoring (2: 2014, 3: 2015, 4: 2016) under the general model of GWASpoly. SNPs with q-values < 0.20, 0.10 and 0.05 are shown in yellow, orange and red, respectively. 5: Local density in highly associated SNPs (p-value < 0.01) for each of the three years of scoring (2014, 2015 and 2016 are shown with blue, green and purple lines). QTLs and metaQTLs from Lopez-Arias et al., 2020 based on three segregating populations (connected by male resistant parent Rosa x wichurana) are also shown. The candidate R genes correspond to an expert annotation of NBS-LRR encoding genes (Lopez-Arias et al., 2020). Results for SNPs located on chr00 (corresponding to all unanchored scaffolds in Hibrand-Saint Oyant et al., 2018) are not shown here (but see Fig. S15 for non-circular Manhattan plots of the p-values including chr00).

The rose GWAS explorer is a tool designed to accelerate research on rose breeding. As a textbook example, we have decided to focus on the identification of genomic regions that correlated with the resistance to the blackspot disease, which is one of the most impactful diseases in garden roses. To identify some regions, we performed GWAS for three years of scoring (Fig. 4, see also Fig. S15). Based on the general model of GWASpoly, the number of significantly associated SNPs greatly varies depending on the year of scoring (red dots, Fig. 4). Additional associated regions are also observed on some other chromosomes for some of the years, including on a region of chromosome 3 that collocates with a meta-QTL recently identified in three different experimentally segregating populations (Lopez Arias *et al.,* 2020). It should be noted that the associated QQplots (Fig. S16) can also highlight an inflation of p-values, which varies in intensity from one model to another, but that can be substantial for the general model (Fig. S16). To facilitate interpretation of the results in this context, we included local enrichments of associated SNPs (*i.e.* number of associated SNPs relative to the total number of SNPs within a 1-Mb window, inner track, Fig. 4). However, this limitation related to p-value inflation should be considered when interpreting the results for this trait, as well as for the other traits displayed on the Rose GWAS browser.

## Discussion

### Evolution of the genetic diversity in roses

Genetic diversity is one fundamental unit of biodiversity since sufficient within-species diversity is needed to face different threats, including diseases, pests, changes in climate, as well as more intrinsic factors associated with evolution under low effective population sizes, and therefore reduce mid- and long-term extinction risks. The adopted post-2020 framework of the Convention on Biological Diversity (CBD, 2022) includes central ambitions regarding the maintenance and the restoration of the genetic diversity in wild species, but also in domesticated species, in order to safeguard their adaptive potential (Goal A & Target 4). It has been recently reported a sharp reduction of the genetic diversity in bread wheat, with a loss of diversity of around 30% over the 20^th^ century (Pont *et al.,* 2019). For ornamentals, for which domestication is far more recent (Purugganan, 2022), little is known about the large-scale genomic impacts of plant breeding.

In the present study, we have questioned how the genetic diversity has evolved during the 19th century, a century corresponding to the golden age for European rose breeding, with an exploding number of varieties (estimated to be around 100 in 1800 and to be 8,000 in 1900; Marriott, 2003, Oghină-Pavie, 2021). Our result is consistent with a substantial reduction of diversity, of around 27.5% in a century in Europe (Fig. 2D), comparing the ancient European and the hybrid tea roses. This reduction has likely taken place across a very limited number of generations (< 10, Fig. 2C, but see also Supplementary Note 2) and is associated with both the backcrossing with a pre-selected Asian genetic background (Figs. 2 & S6-9) and with additional footprints of artificial selection (Fig. 3).

Although this observed reduction of genetic diversity could be considered as more minor regarding a horticultural species as compared to a crop species such as bread wheat, it still holds implications for humans. Indeed, roses represent a common heritage of mankind and serve as a significant cornerstone in the ornamental sector, in addition to being used as a source of food and medicine in some societies. In roses, grafting however makes the long-term maintenance of ancient varieties possible, allowing the backcrossing of modern varieties, improved regarding some specific traits, with more ancient and genetically diverse varieties. This restoration of this genetic diversity is however only possible if the ancient garden roses are maintained over the long term. In roses, two main action targets should be therefore proposed: (i) increasing effort, especially regarding state funding, for safeguarding gardens containing botanical and ancient varieties, and (ii) alert the stakeholders of the importance of maintaining and restoring the genetic diversity in order to ensure a sustainable breeding for long-term, which could be associated with new future breeding targets (*e.g.* new emerging diseases, anthropogenic disturbances or ecosystemic services in urban areas).

### Looking back in time to identify selection targets

Another tremendous advantage of the use of grafted garden roses as domesticated plant models is our ability to phenotype varieties bred during more than a century all together at the same time in a common rose garden. Phenotypically, we indeed have reported important temporal changes in the number of petals, the ability of roses to bloom several times in a year and the susceptibility to the black spot disease (Fig. 1), even if the latter trait evolution was likely unintended. The temporal increase in the number of petals is relatively continuous until 1870, before a slight reduction on average. This is consistent with a change in aesthetic preferences, since (i) varieties with up to 60 petals are nowadays preferred to obtain high-centered buds when the rose flowers are opened, (ii) the rosomania had led to the establishment of rose connoisseur communities from 1880, interested by new targets of selection, including wild-like phenotypes such as single flower (five petals) and spring roses (Oghina-Pavie, 2020). Indeed, at the end of the 19th century, additional rose species were used as breeding materials, such as *R. multiflora* and *R. rugosa*, with single flower and non-recurrent blooming phenotypes. This would be consistent with the apparent progression toward a bimodal distribution for this specific trait and questions the diversification of the consumer tastes at that time, including the return to simple flowers for some varieties (Fig. 1; see also Oghina-Pavie, 2020). Roses of a given period should not be considered as a homogeneous ensemble, they are rather representative of different tastes and breeding objectives among breeders, in such a way that the history of breeding should not be oversimplified. Similarly to petal counts, the susceptibility to black spot has increased until 1890, before a reduction on average, again associated with a marked bimodal distribution (Fig. 1).

The extension of the flowering period by selecting recurrent blooming varieties was often assumed to be the main target of selection during the 19th century. Consistently, during this period, we indeed observed a continuous increase of an index that measures the ability of a rose to rebloom (Fig. 1). Such a continuous selection has likely contributed to generate a marked footprint of selection in rose genomes, since the sharpest signature of the selective sweep is observed above one of the two breaking points (chromosome 3, 24.6-28.2) of a previously characterized 5-Mb inversion of the *RoKSN* gene, *RoKSN^NULL^* allele (chromosome 3, 28.8 - 33.1 Mb, Hibrand-Saint Oyant *et al.,* 2018; Kawamura *et al.,* 2022). However, using the less stringent criteria for detecting sweeps, five additional windows were detected near the second breaking point of the inversion (32.5 - 33.1Mb, see Table S4), therefore providing additional support that this footprint is associated with the artificial selection of Asian alleles at *RoKSN*. Consistent signal was also observed at the second breaking point (Fig. 3), in agreement with the selection in this region. Several alleles at the *RoKSN* gene leading to the extended period were described, including a *RoKSN^copia^* and a *RoKSN^NULL^*. Previously, it was demonstrated a progressive selection of the *RoKSN^copia^* allele during the 19th century (Soufflet-Freslon *et al.,* 2021). This allele is not present in the reference genome we used, this genome contains a large rearrangement at the *RoKSN* locus leading to the *RoKSN^NULL^* allele (Hibrand-Saint Oyant *et al.,* 2018). In the immediate vicinity of the inversion, we find also an homologue of *AP2/TOE* (RC3G0243000, 33.2Mb), another crucial gene controlling the double flower phenotype and therefore partly the number of petals (François *et al.,* 2018; Hibrand-Saint Oyant *et al*., 2018). However, *AP2/TOE* is unlikely to be the gene responsible for this footprint (see Supplementary Note 1), suggesting that recurrent-flowering alleles of RoKSN were the main targets of artificial selection. This rearrangement may explain the complexity of the local footprints associated with the nucleotide diversity and Tajima’s D landscapes. Additional footprints have been also uncovered, but currently were not associated with any previously reported genes. In the future, thanks to more numerous whole-genome sequences available, as well as the new technological and methodological developments allowing the access to phased polyploid genomes, a much clearer picture could be anticipated. Extending these investigations to the 20th century could be of particular interest since more modern roses are characterized by a clearer distinction between the genetics used for garden *vs.* cut roses. Cut roses - especially red ones - have potentially experienced a more intense reduction of global genetic diversity and more pronounced local footprints of selection, as typical expected based on the huge difference of the two respective markets (*e.g.* in France in 2018, garden roses: 50M€, cut roses: 376M€, Clotault *et al*., 2022).

### Advances and challenges for GWAS in rose

As a tool for the future of rose breeding, we have released the largest GWAS catalog for roses (https://roseGWASbrowser.github.io/). Our GWAS browser allows the website visitors to explore the association with more than 25 traits, on the basis of our panel of 204 accessions. The objective is to make future breeding more effective for some traits, including the resistance to the black spot disease (Fig. 4). This trait represents a typical example of an important target need in order to find sources of genetic resistances in roses, in the context of a tightening environmental legislation associated with the use of chemicals due to their deleterious environmental impacts and health risks.

Despite the great advance made with the release of the GWAS catalog, the direct insights appear relatively limited to date. Indeed, the GWAS strategy in our panel has generated few repeatable and easily interpretable signatures of associations. Overall, the relative lack of resolution is likely associated with the difficulty of performing GWA studies in populations for which phenotypes still highly correlate with the population structure gradient as a result of the limited number of generations (*e.g.* see Fig. S4). Indeed, similar results were observed for most of the investigated traits. As a consequence, all the results should be interpreted with some caution, requiring further validation through independent analyses. Regarding the black score disease, some potential genomic regions of interest were detected, including a genomic region at the very end of chromosome 4 that is associated with the three years of scoring (Fig. 4). Additional locations are also observed on chromosome 3 for some of the years, including in a region that collocates with a meta-QTL recently identified in three different experimentally segregating populations, which were phenotyped for black spot disease over several years and locations (Lopez Arias *et al.,* 2020).

As compared with the use of QTL mapping in experimental crosses, extensive GWAS analyses seem to have a more limited benefit in rose than on some other domesticated or non-domesticated species (*e.g.* Korte & Farlow, 2013; Alqudah *et al.,* 2020). Several non-mutually exclusive reasons could explain this paradox: (i) a too limited number of varieties used in our study, (ii) the limited number of generations of recombination in the investigated collection (Fig. 2C) that could contribute to maintaining a predominant association with the population structure, leading to a low statistical power in the model. Combining GWAS and QTL approaches (Fig. 4), when available, offers the possibility to increase the statistical support for some regions of interest, extends the allelic diversity by considering other alleles than those segregating between the parents, describes the genomic locations of the associated regions and prioritizes the research effort. As a typical example, the last third of chromosome 3 could appear as a more promising region for selection, therefore guiding the research efforts in this direction. Remarkably, chromosome 3 has been associated with many traits in rose (resistance to black spot, but also number of petals, fragrance, prickles, recurrent blooming, self-incompatibility; *e.g.* Hibrand-Saint Oyant *et al.,* 2018, Kawamura *et al.,* 2022), which could therefore suggests the presence of tradeoffs between some traits of interest. A more detailed genomic characterization of the genomic landscapes, including local linkage disequilibrium patterns, based on more modern varieties, is especially needed on this chromosome.

## Conclusion

Combining large phenotypic data and genetic investigations using 204 individuals genotyped using a high density SNP array, plus 32 whole-genome sequences, we found support for both phenotypic and genomic changes during the 19th century. With an unprecedented accuracy, we confirmed the progressive shift from a European to a more Asian-like genetic background, with Hybrid tea roses exhibiting a three-quarter Asian genetic background. This genetic makeup likely corresponds to several generations of backcrossing of the early European x Asian varieties with the ancient genetic background, as it was previously suggested (Liorzou *et al.,* 2016, see also Supplementary Note 3). As a result of the lower resident diversity in the ancient Asian varieties, partly due to a more ancient and intense history of selection in Asia, as well as to some specific genomic footprints of artificial selection - especially in the region of *RoKSN -*, rose breeding in Europe has contributed to a substantial loss of genetic diversity during the 19th century (up to 27.5% in hybrid tea roses). In addition to providing a fine description of the history of rose breeding, we have generated the largest GWAS catalog to date, with the aim of providing fundamental knowledge for the future of rose breeding.

## Acknowledgments

This present article is dedicated to the memory of our cherished colleague, Laurence Hibrand-Saint Oyant, who passed away in November 2024 (see our dedication section, Supporting Information). This article has been peer-reviewed and recommended by Peer Community In Evolutionary Biology (https://doi.org/10.24072/pci.evolbiol.100793). We are grateful to our PCI recommender Mathieu Gautier and the three reviewers (Pierre Nouhaud, Vincent Segura and an anonymous reviewer) for having provided detailed and constructive reviews. We acknowledge the Roseraie Loubert (Gennes-Val-de-Loire, France) and Biological Resource Center Pome Fruits and Roses (Angers, France) for maintaining the genetic resources and their associated data. We also acknowledge the Gentyane facility (INRAE Clermont-Ferrand, France) and the ANAN platform of the SFR Quasav (Angers, France; including Muriel Bahut) for the genotyping data. We would like to thank the Next Generation Sequencing Facility at Vienna BioCenter Core Facilities (VBCF), member of the Vienna BioCenter (VBC), Austria. We are grateful to the genotoul bioinformatics platform Toulouse Occitanie (Bioinfo Genotoul, https://doi.org/10.15454/1.5572369328961167E12) for providing computing and storage resources. We also thank Jérôme Chené (Pépinières Roses Loubert, Gennes-Val-de-Loire, Pays-de-la-Loire, France) for providing fresh plant material of a few whole-genome resequenced individuals. We also thank Roeland Voorrips and Giorgio Tumino for their valuable advice regarding fitPoly and GWASpoly at the beginning of the project.

## Author contributions

T.Le., E.A., S.B., J-C.C., C.O.P, F.F., L.H.S.O & J.Cl. designed research; T.Le., E.A., T.T., A.C, J.Ch, T.Lo., A.P., V.S.F, F.F., L.H.S.O & J.Cl contributed to the sampling; A.C. & L.H.S.O. performed DNA extractions; J.J. performed the *RoKSN* and *RoAP2 genes* genotyping; T.T., J.C., T.Lo., A.P. & V.S.F contributed to the phenotyping; S.B., J-C.C. & S.N.P. contributed to the GC-mass spectrometry data; T.Le & E.A. analyzed data under the supervision of C.O.P, F.F., L.H.S.O. & J.Cl.; T.Le. drafted the manuscript. All authors have subsequently read, edited and approved the manuscript.

## Data availability

Raw sequencing data of the newly sequenced roses are publicly available on the Sequence Read Archive (SRA) under the BioProject PRJNA997103. Scripts were made available on Zenodo (https://doi.org/10.5281/zenodo.14450241). The GWAS rose browser is available at https://roseGWASbrowser.github.io.

