## Supporting Information for "Dark side of the honeymoon: reconstructing the Asian x European rose breeding history through the lens of genomics"

Peer-reviewed & recommended by PCI Evol Biol

#### Supporting Information

**Dedication** **p. 2**

**Supplementary Notes** **p. 3-13**
*Sup. Note 1* 3
*Sup. Note 2* 7
*Sup. Note 3* 11

**Supplementary Figures** **p. 14-29**
*Figure S1* 14
*Figure S2* 15
*Figure S3* 16
*Figure S4* 17
*Figure S5* 18
*Figure S6* 19
*Figure S7* 20
*Figure S8* 21
*Figure S9* 22
*Figure S10* 23
*Figure S11* 24
*Figure S12* 25
*Figure S13* 26
*Figure S14* 27
*Figure S15* 28
*Figure S16* 29

**Supplementary Tables** **p. 30-41**
*Table S1* 30-31
*Table S2* 32-36
*Table S3* 37
*Table S4* 38-41

### Dedication

This present study is dedicated to the memory of our colleague, Laurence Hibrand-Saint Oyant, one of the two senior authors of this work. Laurence unexpectedly passed away on November 11, 2024. Her contributions to this study were invaluable, encompassing everything from its design to the interpretation of the results.

Beyond her pivotal role in this research, Laurence was a key figure in the field of rose genetics and genomics, as typically shown by the sequencing of one of the two first high-quality rose genomes (*e.g.* Hibrand *et al.*, 2018 *Nature Plants*). Her scientific legacy has profoundly shaped our understanding of rose breeding and will continue to inspire researchers in the field for years to come. Laurence was deeply committed to mentoring young scientists, and she defended the place of women in science, a cause that was close to her heart.

Laurence was not only a brilliant scientist, but also a cherished colleague whose positivity and warm smile left an indelible mark on everyone she worked with. She was deeply admired and appreciated at the Institute of Research in Horticulture and Seeds (IRHS) in Angers, France. Her loss is felt deeply by all who had the privilege to know her, both professionally and personally. Our deepest thoughts are with her husband Alain, her two children, Paul and Arthur, and all those who had the chance to interact with her, in both her professional and personal life.

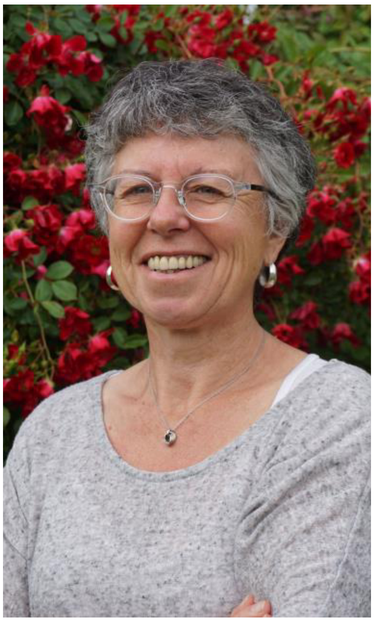
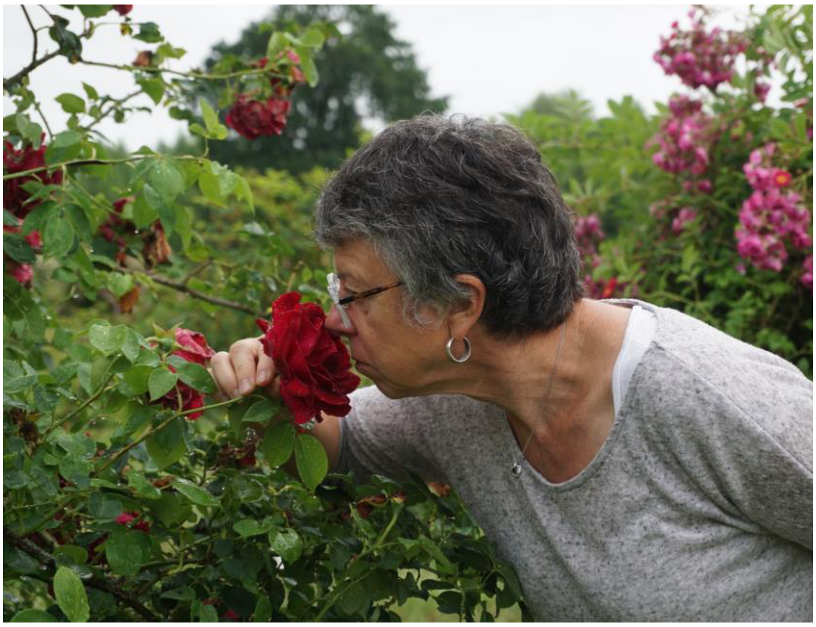

*Laurence Hibrand-Saint Oyant, our late colleague, pictured at the Loubert ancient rose garden in the Loire Valley, France, where most of the samples for the present study were collected (pictures by N. Mansion, INRAE).*

### Supplementary Note 1

On chromosome 3, two loci controlling two important ornamental traits are close to each other and located in the vicinity of a region exhibiting footprints of artificial selection (see Fig. 3). To go further in the analysis, we have genotyped the 204 genotypes of GWAS for different known alleles of these two genes: *RoAP2* (controlling double flower) and *RoKSN* (controlling recurrent blooming).

#### ***RoKSN* alleles**

We have previously demonstrated that the continuous flowering phenotype is due to the mutation of floral repressor, a rose *TERMINAL FLOWER1* homologue, denominated *RoKSN* (Iwata *et al.*, 2012). Specifically, we have identified an allele leading to continuous flowering. The insertion of a *copia* transposable element in the *2nd* intron of *RoKSN* leads to the absence of *RoKSN* transcript accumulation (*RoKSN$^{copia}$* allele). Recently, by sequencing *R. chinensis* 'Old Blush', we have detected a large rearrangement on the chromosome 3 leading to the *RoKSN* deletion, *RoKSN$^{null}$* allele (Hibrand Saint-Oyant *et al.*, 2018). By sequencing different garden roses, a new allele was detected: at position 181 of the *RoKSN* coding sequence a G is converted into a A. At the homozygous stage, this *RoKSN$^{181A}$* allele was shown to be associated with a weak capacity to rebloom. This change is not affected the activity of the protein, but less *RoKSN* transcripts are accumulated (Soufflet-Freslon *et al.*, 2021).

Using different primers, we have genotyped all these alleles based on previous described methods for *RoKSN$^{copia}$* (Iwata *et al.*, 2012), *RoKSN$^{null}$* (Kawamura *et al.*, 2022), *RoKSN$^{A181}$ and RoKSN$^{G181}$* (Soufflet-Freslon *et al.*, 2021). On the garden rose collection from the 19$^{th}$ century, we have detected 11 different combinations (Supplementary Note 1 Fig S1). Since we used a genotyping by PCR strategy, we are unable to dose the different alleles into polyploid roses and consequently, we only noted the presence or absence of the alleles.

To quantitatively phenotype the ability of a rose to rebloom, we have scored the number of roses on each week for 3 years from spring to autumn. We have previously developed a statistical method on the flowering curve using Gaussian mixture models and generated indicators, such as the Reb.Mag parameter for the magnitude of reblooming (Proïa *et al.*, 2016). A value of 0 means that the roses flower only in spring (once-flowering roses), the higher the value (above 0), the higher the reblooming capacity.

Comparing genotype and phenotype data, we observed a large variability in the reblooming ability of the different garden roses (Supplementary Note 1 Fig. S1). The roses with the *RoKSN$^{copia}$* allele have the highest ability to rebloom (median of 0.42). The roses with the *RoKSN$^{null}$* allele also exhibit a high index, albeit lower than those with the *RoKSN$^{copia}$* allele (median of 0.21). These two alleles were described to bring continuous flowering at the homozygous stage (Iwata *et al.*, 2012; Hibrand Saint-Oyant *et al.*, 2018). The reblooming ability of the roses without the *RoKSN$^{copia}$* or *RoKSN$^{null}$* alleles is very weak with the majority of these roses with a Reb.Mag value of 0 (once-flowering roses). As previously demonstrated, the roses with the *RoKSN$^{A181}$* allele have a weak ability to rebloom (Soufflet-Freslon *et al.*, 2021). The heterozygous roses (*RoKSN$^{A181}$* or *RoKSN$^{G181}$ with RoKSN$^{copia}$* or *RoKSN$^{null}$*) have an intermediate phenotype with a weak ability to rebloom, suggesting that the *RoKSN$^{G181}$* is not completely dominant or other loci control the ability to rebloom as previously suggested (Soufflet-Freslon *et al.,* 2021). Interestingly the combination of *RoKSN$^{A181}$* and the *RoKSN$^{copia}$ or roKSN$^{null}$* allele leads to rose with a higher, albeit non-significant, ability to rebloom when compared with the *RoKSNG181* allele (Supplementary Note 1 Fig. S1).

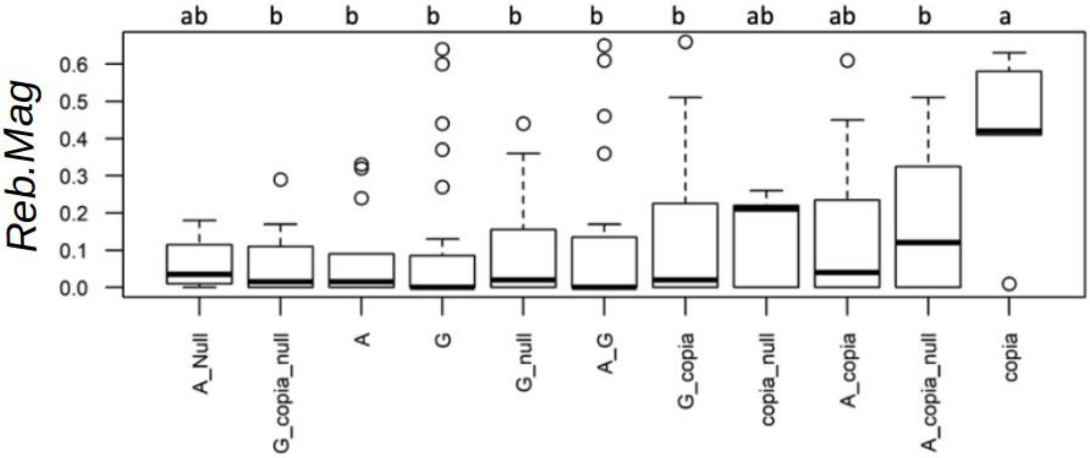

*Supplementary Note 1 Figure 1: Phenotypic variation for the recurrent blooming estimated with the Reb.Mag index (Proïa et al., 2016) according to the genotypes at the RoKSN gene in garden roses from the 19$^{th}$ century. Average value for Reb.Mag among the 3 years of scoring is presented. A value of 0 for Reb.Mag means that the roses are once-flowering roses). Higher is the value, higher is the magnitude of reblooming. The 11 different allelic combinations at the RoKSN locus are presented. A is corresponding to the RoKSN$^{A181}$ allele, G to RoKSN$^{G181}$, null to RoKSN$^{null}$ and copia to RoKSN$^{copia}$. Only the presence of the allele is scored (no dosage for polyploid roses).*

**<u>RoAP2 alleles</u>**

Previous studies have demonstrated the important role of a rose *APETALA2/TOE* homologue (*RoAP2*) in the control of a double flower in roses (François *et al.*, 2018; Gattolin *et al.*, 2018; Hibrand-Saint Oyant *et al.*, 2018), however this gene is unlikely the reason of the main footprint identified on chromosome 3. In double flowers, the insertion of a *gypsy* retrotransposon element in the 8$^{th}$ exon of the rose *APETALA2* homologue leads to the obtention of a miRNA *RoAP2* resistant allele (Gattolin *et al.*, 2018). This allele has been described to be dominant. We have used a PCR based method to genotype the two alleles: allele WT (no insertion of the retrotransposon) and the allele *gypsy* (insertion of the retrotransposon). Roses bearing the *gypsy* alleles are supposed to have a double flower phenotype (François *et al.,* 2018; Gattolin *et al.,* 2018; Hibrand Saint-Oyant *et al.,* 2018). Similarly than for *RoKSN*, it should be noticed that the dosage of the allele in polyploid plants was not possible using this genotyping method.

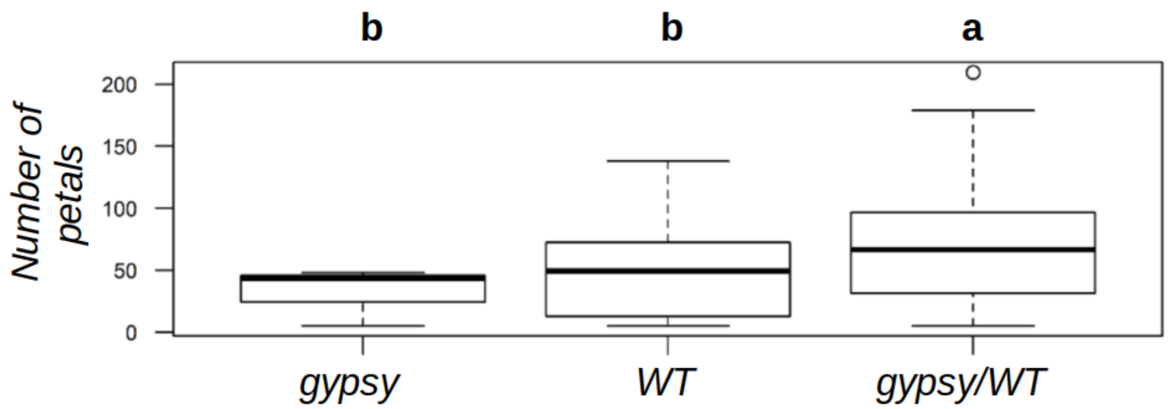

*Supplementary Note Figure S2: Phenotypic variation for the number of petals according to the genotypes at the RoAP2 gene in garden roses from the 19th century. The recessive WT allele corresponds to the allele with no gypsy retrotransposon. The dominant gypsy allele corresponds to the miRNA-resistant allele with a gypsy retrotransposon insertion.*

In the garden rose collection from the 19th century, roses bearing only the WT alleles are largely double (median with 49 petals), even if roses with the *gypsy* allele present significantly more petals (median with 67 petals, Supplementary Note Fig. S2). These results are not in agreement with previous results showing that roses with only the *WT* allele present simple flowers. These previous results are mainly based on only a few modern roses (François *et al.*, 2018; Gattolin *et al.*, 2018) or a large GWAS panel from the 20th century (Hibrand Saint-Oyant *et al.*, 2018). This discrepancy between the previous results and our results could be explained by the fact that we are working with ancient material (roses from the 19th century), and could suggest a different genetic determinism between roses from the 19th and 20th centuries. Double flower roses already existed in old European and Chinese roses, with different genetic origins, which could therefore suggest that other loci may be present in the garden roses from the 19th century. Additional research on the genetic bases of the number of petals in rose is required, potentially focusing on other associated regions (https://rosegwasbrowser.github.io/numberofpetals/).

**Conclusion**

Despite that we cannot completely rule out the hypothesis of a third gene that could have contributed to the footprint observed in Fig. 3 (especially given the length of the region), our results are far more consistent with the hypothesis of an artificial selection of *RoKSN* alleles for an extension of the blooming period than an hypothesis involving the AP2 gene. We therefore suggest that the intense selection for an extended blooming period (Fig. 1B) explains the genomic footprints observed in Fig. 3. This selection could be associated with the selection of roses with *RoKSN*$^{copia}$ or *RoKSN*$^{null}$ alleles. Such a selection has already been demonstrated for the *RoKSN*$^{copia}$ allele (Soufflet-Freslon *et al.*, 2021).

In addition, it should be noted that large among-variety variances are observed in the ability to rebloom among genotypes with the same combination of alleles at *ROKSN* (Supporting Note 1 Fig. 1), suggesting a more complex genetic basis for this trait, with additional genes contributing to this phenotypic variation.

### Supplementary Note 2

#### Introduction

Cultivated garden roses exhibit a broad range of ploidy levels, with roses at the diploid, triploid (3x = 21), and tetraploid (4x = 28) levels, and some even higher. As part of a former study, we investigated population structure at 32 microsatellite markers across more than 1,200 varieties (Liorzou *et al.*, 2016), providing an initial estimate of probable ploidy levels based on the observed allele count, despite the limitations associated with this type of marker (*e.g.* overlapping patterns, stutter artifacts).

#### Strategy used

Given that at the time of sampling (first sampling campaigns in 2020, followed by a second campaign in 2021), we already knew that genotyping would be conducted using the 'WagRhSNP' Axiom rose genotyping array, which was developed specifically for tetraploid garden and cut roses, we decided to preferentially collect samples from both diploid and tetraploid roses. Importantly, it was not feasible to conduct this project solely with tetraploid roses, since most ancient Asian roses were presumed to be diploid. Our hypothesis was that diploid species would mostly be genotyped as AABB on the array when diploid and tetraploid samples are genotyped together. Subsequently, the same strategy of sampling both diploid and tetraploid roses was applied for the sequencing phase of our project (sampling campaign in 2021). For sequencing, we were able to fully account for the ploidy level to generate unbiased estimates of nucleotide diversity, a core objective of the project. Concretely, we first inferred the most likely ploidy level for each individual based on a first round of SNP calling. We then assumed this inferred ploidy level to genotype individuals at a variable ploidy level in a second round of SNP calling.

#### Accuracy of the ploidy inference for the sequenced individuals

Importantly, coverage is highly variable among the different sequenced individuals. This can have an impact on the accuracy of our inferences. To provide a more precise view on the variation of the depth of coverage among the sequenced individuals, we summarized the values at 5% of all SNP positions for the 32 individuals (DP field, Table S2). The coverage greatly varies, from 5.2 (20_Victor_verdier) to 44.5 (04_Rosa_gallica_officinalis). Aware of the potential impact of low coverage on the quality of the ploidy inference, we decided to estimate allelic balance for the minor allele only for SNP that has a minimum coverage of 20. However, this strategy still has two limitations: i) the number of SNPs meeting the minimum coverage threshold of 20, which are used to generate the allelic balance distributions, differs significantly in magnitude, and ii) the precision of the allelic balance estimates varies greatly depending on the coverage (see: Supplementary Note 2 Fig. 1). Typically “04_Rosa_gallica_officinalis” (mean_cov=44.5) and “19_Yellow_island” (mean_cov=33.9) exhibit near perfect tetraploid profiles (see Fig. S2, Supplementary Note 2 Fig. 1 and “Extended_FigS2_ploidy_chr_per_chr.pdf” in the Zenodo repository for details), while other tetraploid individuals exhibit much lower genome-wide coverage (all tetraploids: 5.2 - 27.3, mean=10.6, non-Botanical tetraploid accessions: 5.2 - 15.7, mean= 8.3), penalizing the quality of the local model fitting (loess).

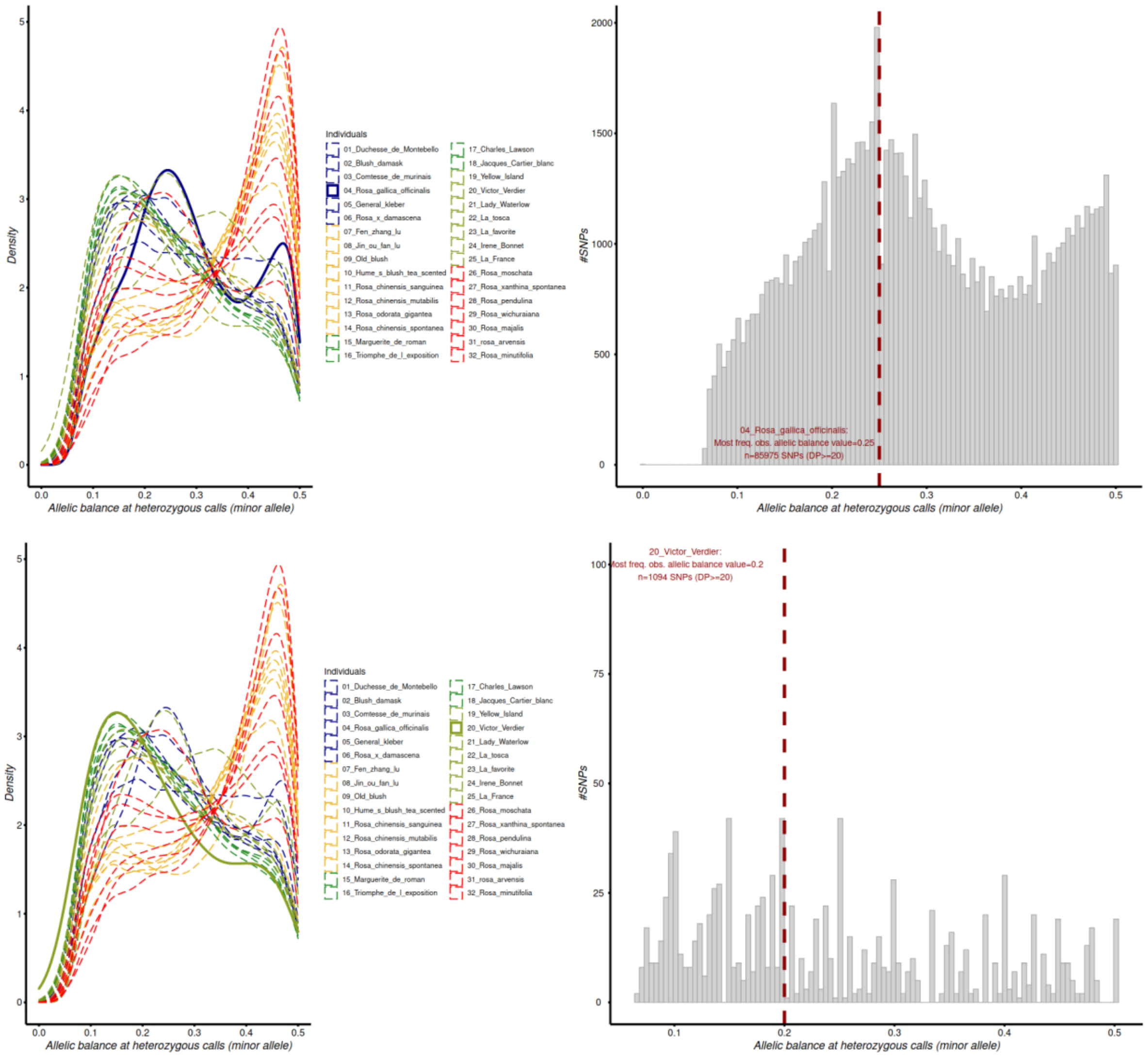

*Supplementary Note 2 Fig. 1: DIfference of accuracy in identifying the ploidy level depending on the depth of coverage. This figure highlights the most contrasted individuals in the dataset regarding the coverage, with Rosa gallica officinalis (top, mean_cov=44.5) and Victor Verdier (bottom, mean_cov=5.2). Most frequently observed allelic balance corresponds to the exact allelic balance among the SNPs with DP>20. Deviations from 0.25 for individuals with low coverage should not be overinterpreted given the high stochasticity in the distributions. Results for all the other individuals are available on the Zenodo repository (“*Extended_FigS2_ploidy_chr_per_chr.pdf”).

Nevertheless, for all except two individuals, the most frequently observed allelic balance value among all values corresponds to 0.25, 0.33 or 0.5 (see “script_infer_ploidy_from_data.R” and “Extended_FigS2_ploidy_chr_per_chr.pdf” in the Zenodo repository for details), consistent with expectations for tetraploids, triploids and diploids, respectively. The only two exceptions are “20_Victor_Verdier*”* and “21_Lady_Waterlow” (0.2), but this does not necessarily indicate that these individuals are pentaploids. These two also have the lowest number of SNPs meeting the coverage threshold of >20, where allelic balance was applied, further complicating the context for accurate ploidy inference (see Supplementary Note 2 Fig. 1). For these two individuals, the tetraploidy we considered based on the distributions of allelic balance is consistent with the results of Mathilde Liorzou and collaborators (2016).

#### Accuracy of the population structure inference

Given that the PCA we reported in Fig. 2B assumes diploid calls from the initial SNP calling round, we investigated the potential impacts of non-strict diploidy in the data. Concretely, we generated an allele count matrix from the AD field of the VCF using the vast majority of our biallelic SNPs (65 million SNPs). The treemix-formatted input file was subsequently imported in the R package 'poolfstat' thanks to the genotreemix2countdata function in order to perform a random allele PCA under 'poolfstat'. The advantage of this hypothesis is that it does not rely at all on the diploid calls, but is just based on the allele counts. We observed that the results are highly consistent (Supplementary Note 2 Fig. 2), with extremely minor variation of both the proportions of explained variance and the relative locations of the individuals on the different PCs, suggesting that our population structure is highly robust to the variable ploidy level among individuals.

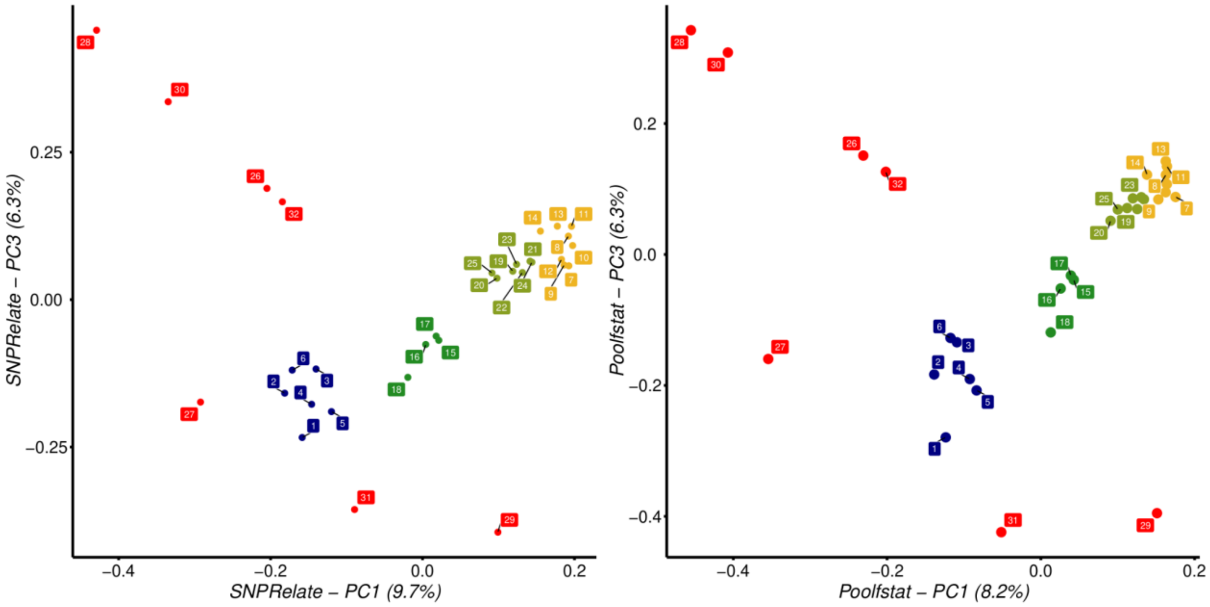

*Supplementary Note 2 Fig. 2: Comparison of the results of PCA assuming the diploid calls (Fig. 2B, left) or directly based on the allele counts (random allele PCA from poolfstat, right) for the first and third components.*

#### Accuracy of the KING inference

KING was developed for diploid species. Since KING analysis was conducted on the initial SNP calling round, resulting in diploid calls, we were able to use KING in our SNP set, however we cannot rule out a potential impact of variable ploidy levels among the roses on the behavior of the software. Given that most samples identified as family related are tetraploid (see results), we can have two hypotheses. First, that the analysis performed under KING is robust to this bias and indeed recovered more family relationships in tetraploids, which could indeed be expected since modern roses are mostly tetraploids. Second, assuming that this result is partly associated with bias in the kinship inference, it is important to note that this result would therefore be in the direction of more easily identifying relationships in tetraploids. Given that our main objective was to exclude the most related individuals for all the subsequent analyses to ensure accurate results (*e.g.* nucleotide diversity), it would suggest that our downstream analyses are even more conservative for the three groups only composed of tetraploids (*i.e.*

ancient European, early Asian x European and hybrid tea roses). However, this limitation associated with deviation from true diploidy should be borne in mind when interpreting the results in terms of the number of generations of intercrossing.

#### Conclusion

From our sampling to campaigns to our final analyses, we have made considerable effort to account as much as possible for the variable ploidy in garden roses and that our study also contributes to move forward by solving many challenges and providing ploidy-aware analyses (*e.g.* nucleotide diversity). However, we also recognize that our work did not solve all challenges. Consequently, additional studies will be needed to confirm our results. It is also important to note that polyploidy in roses is even more complex than just considering different number of chromosomes sets, since some works have for instance reported evidence for segmental allopolyploidy in some tetraploid roses (see Koning-Boucoiran *et al.*, 2012; Bourke *et al.*, 2017; Cheng *et al.*, 2024), which further complicates the story. We encourage more work in this direction.

#### Supplementary Note 3

At diagnostic loci between ancient European and Asian samples (allele frequency of 0 and 1 for the reference allele, respectively; see Materials & Methods and Fig. S8), we observed allele frequencies that are roughly similar to 1:1 and 3:1 contributions between Asian and European samples in the Early European x Asian and Ancient Asian groups. Such simple ratios could suggest that our sampling follows some simple Mendel's laws, considering that all Early European x Asian are F1 between Ancient European and Ancient Asian, and that they were subsequently backcrossed with the Ancient Asian group to generate all the hybrid Tea roses used in our study (hypothesis 1, Supplementary Note 2 Figure).

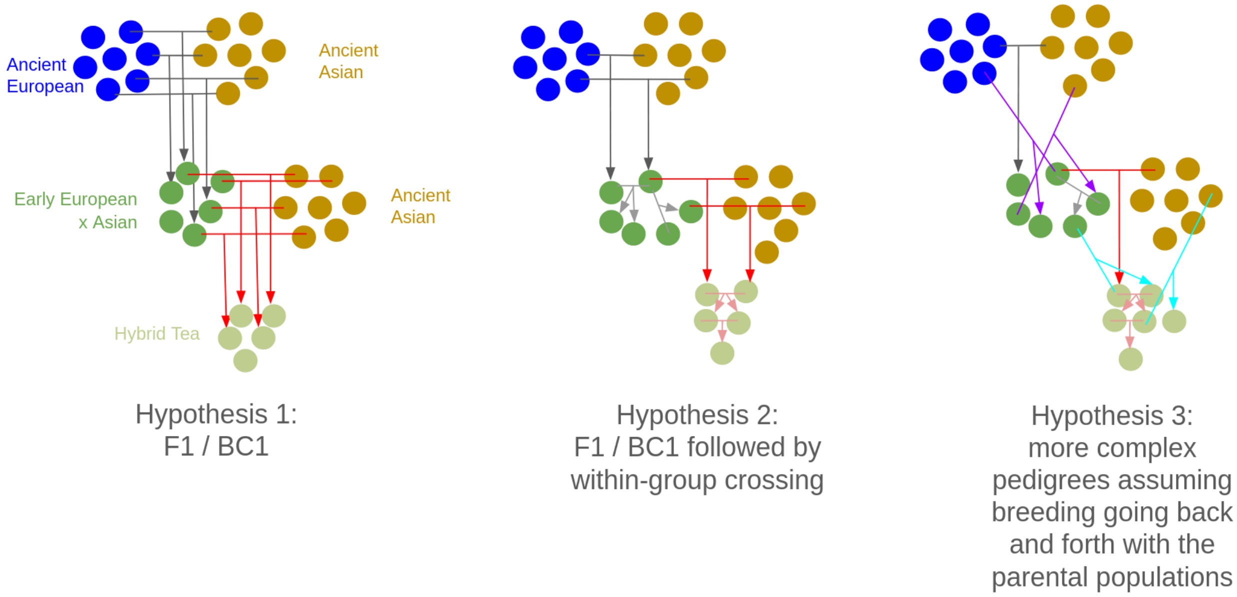

*Supplementary Note 2 Figure : Schematic illustration of three different scenarios that could have led to the overall observed allele frequencies at diagnostic markers (1:1 and 3:1 ratios, see main text). Hypothesis 1 (left) assumes that all Early European x Asian are first-generation hybrids (F1, black arrows) and, similarly, that Hybrid Tea roses are first-generation backcrosses (BC1, red arrows) of the Early European x Asian with the ancient Asian roses. Hypothesis 2 assumes that the Early European x Asian and Hybrid Tea groups initially derived from F1 and BC1, but within-group crosses were possible (grey and pink arrows). Hypothesis 3 (right) relaxes this hypothesis and assumes that artificial selection was less directional than assumed under the two previous hypotheses, allowing among-group crosses (purple and blue arrows).*

To investigate whether all our Early European x Asian are consistent with hypothesis 1, we randomly subsampled several hundreds of diagnostic SNPs covering the whole genome and investigated whether the genotypes are all heterozygous at these locus. Let's assume that all the groups share the same ploidy level (e.g. diploid). Given that the loci are by definition homozygous for the reference and alternate alleles in the ancient Asian and European groups, early European x Asian are expected to be heterozygous (0/1) at all diagnostic loci. Here, the variable ploidy level between groups complicates this expectation, since ancient Asian samples are mostly diploid, while ancient European roses are mostly tetraploid. Nonetheless, assuming the F1 hypothesis, no homozygous calls are expected (0/0/0/0 or 1/1/1/1 for tetraploid). Contrary to this expectation, we instead observe an important proportion of parental genomes (between 58 and 67% of all genotypes, Supplementary Note 2 Table). As a consequence, our results are therefore not consistent with the first hypothesis.

*Supplementary Note 2 Table : observed frequency of the 5 possible genotypes in tetraploids observed at 709 randomly selected diagnostic loci covering the whole genome (from Chr01 to Chr07 only), with allele 0 corresponding to the allele of the reference genome (Old Blush, an Asian genotype). All early European x Asian and hybrid Tea roses investigated are tetraploids.*

| VarietyID | Marguerite_de_Roman | Triomphe_de_lexpo | Charles_Lawson | Jacques_Cartier_blanc | Lady_Waterlow | La_Tosca | La_Favorite | Irene_Bonnet |
|---|---|---|---|---|---|---|---|---|
| SeqID | 175586 | 175587 | 175623 | 175624 | 175626 | 175627 | 175628 | 177143 |
| Group | Early European x Asian | Early European x Asian | Early European x Asian | Early European x Asian | Hyb Tea roses | Hyb Tea roses | Hyb Tea roses | Hyb Tea roses |
| 0/0/0/0 | 0.363 | 0.309 | 0.347 | 0.250 | 0.792 | 0.711 | 0.669 | 0.651 |
| 0/0/0/1 | 0.135 | 0.090 | 0.140 | 0.077 | 0.073 | 0.098 | 0.103 | 0.149 |
| 0/0/1/1 | 0.113 | 0.157 | 0.173 | 0.143 | 0.073 | 0.082 | 0.110 | 0.099 |
| 0/1/1/1 | 0.094 | 0.156 | 0.107 | 0.110 | 0.034 | 0.042 | 0.037 | 0.044 |
| 1/1/1/1 | 0.295 | 0.288 | 0.233 | 0.420 | 0.027 | 0.068 | 0.081 | 0.058 |
| f(1) | 0.456 | 0.505 | 0.435 | 0.594 | 0.107 | 0.164 | 0.189 | 0.177 |

Then, two scenarios can support the observed results similarly well. The first alternative scenario (2nd hypothesis) assumes that all the early European x Asian varieties derive from some first-generation hybrids, which were then subsequently crossed together, maintaining the 1:1 ratio of allele frequency at diagnostic alleles but allowing different classes of genotypes at diagnostic SNPs as observed in the Supplementary Note 2 Table. Under the last scenario (3rd hypothesis), the history of selection is less directional, with crossing going back and forth with the Ancient Asian and European roses. Following this scenario, the observed ratios are indeed close to the textbook examples of Mendel's ratios, but these values only correlate and a causative link would be spurious, following the old adage "*Cum hoc ergo propter hoc*". According to us, the latter hypothesis is the most likely, since rose breeders' interests vary through space and time in such a way that the favorite progenitors are expected to have changed, among breeders and through time, leading to more complex pedigrees than the simple ones depicted in the two first hypotheses. It is also worth noting that rose crosses through hand pollination have not been controlled before 1870, and more broadly, the late 19th century (Oghina-Pavie, 2020, p.75), explaining why such more complex pedigrees are also particularly expected.

Further research aiming at reconstructing the pedigrees based on a large number of varieties will probably help to provide a new complete view about the history of rose breeding. This will be only possible if a considerable effort is made to maintain old rose collections.

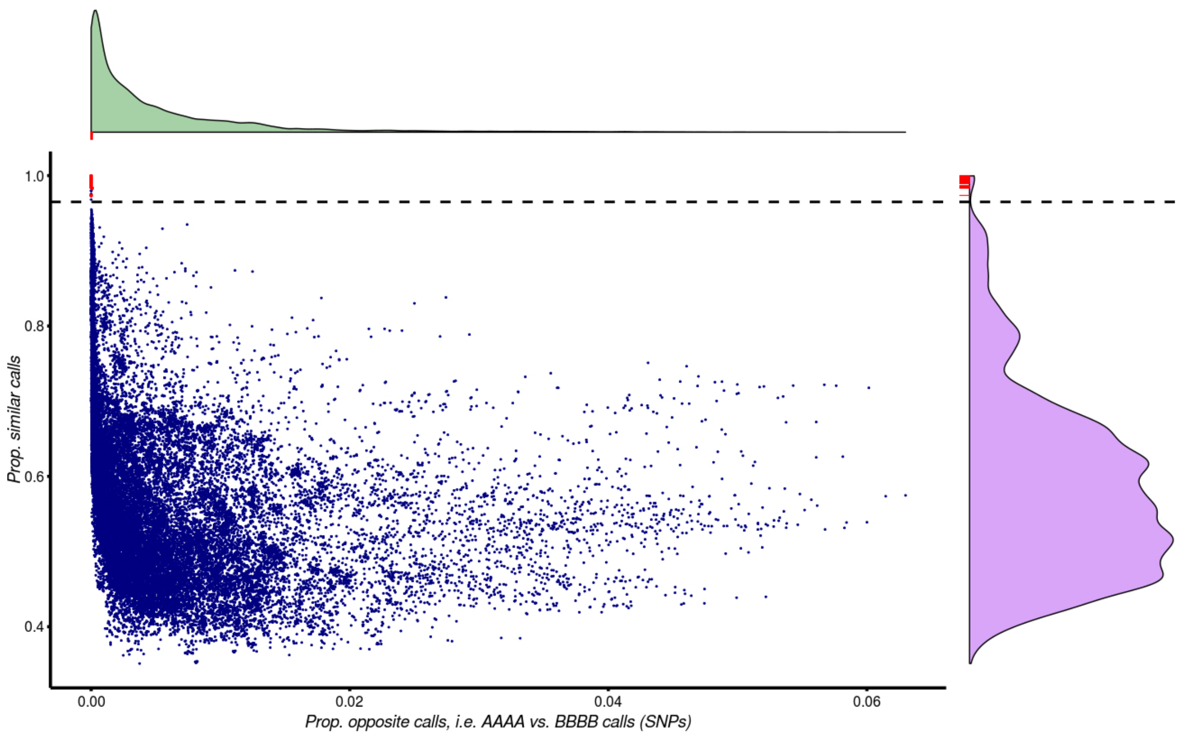

Figure S1: Clone detection in the dataset. Each dot shows the proportion of opposite calls (AAAA vs. BBBB, x-axis) and the proportion of exactly similar calls (y-axis) for all pairs of samples. True replicates of the analyses are shown in red. Distributions of the two metrics are shown. As the proportion of similar calls is more discriminating, we considered the threshold of 96.5% (dotted line) as the minimum threshold for defining a clone. All true replicates included on our dataset are above this threshold (red lines on the purple distribution). See Table S1 for the list of potential clones.

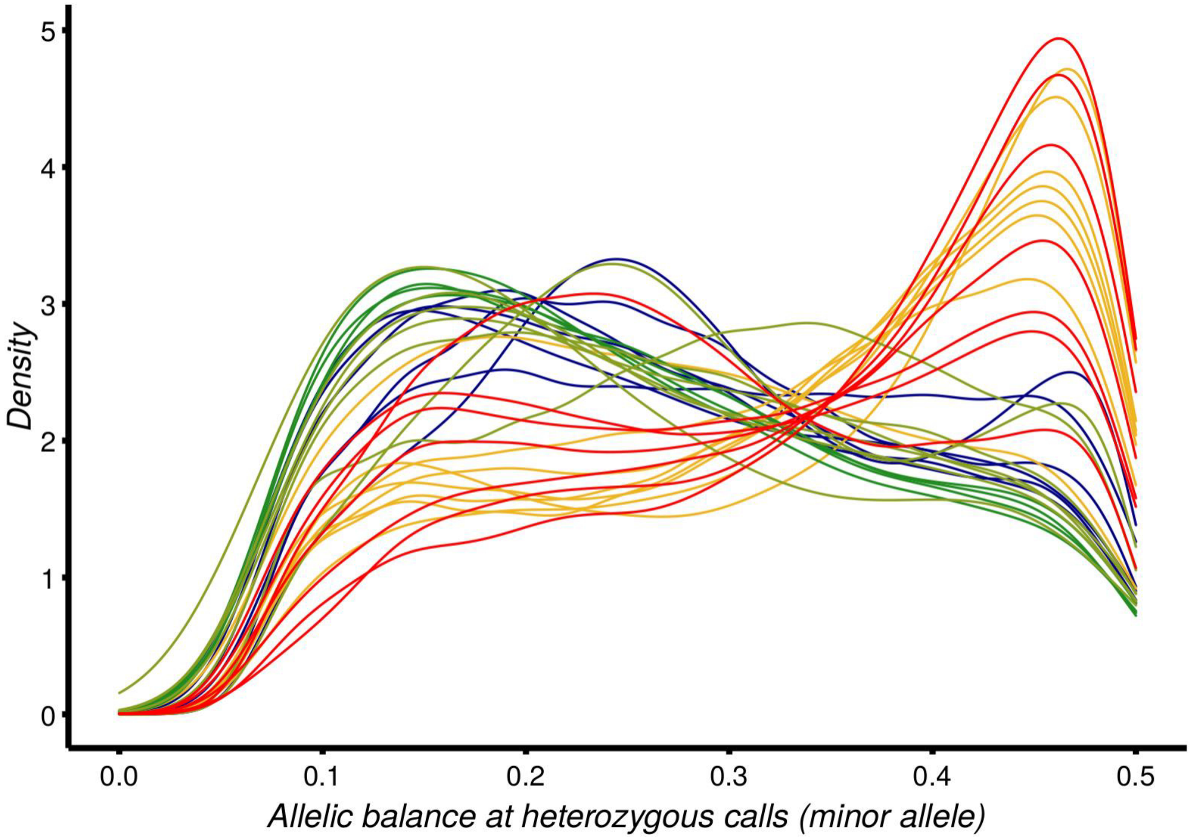

*Figure S2: Inference of the ploidy based on the allelic balance at heterozygous sites. Diploid individuals are expected to have its highest peak near 0.5. Note that given the low to moderate depth of coverage and the fact that we use the allelic balance of the minor alleles, the mode of the distribution is expected to be shifted toward slightly lower values than 0.5 and 0.25 for diploid and tetraploid individuals, respectively. Detail information for each individual is available on the Zenodo repository ("Extended_FigS2_ploidy_chr_per_chr.pdf")*

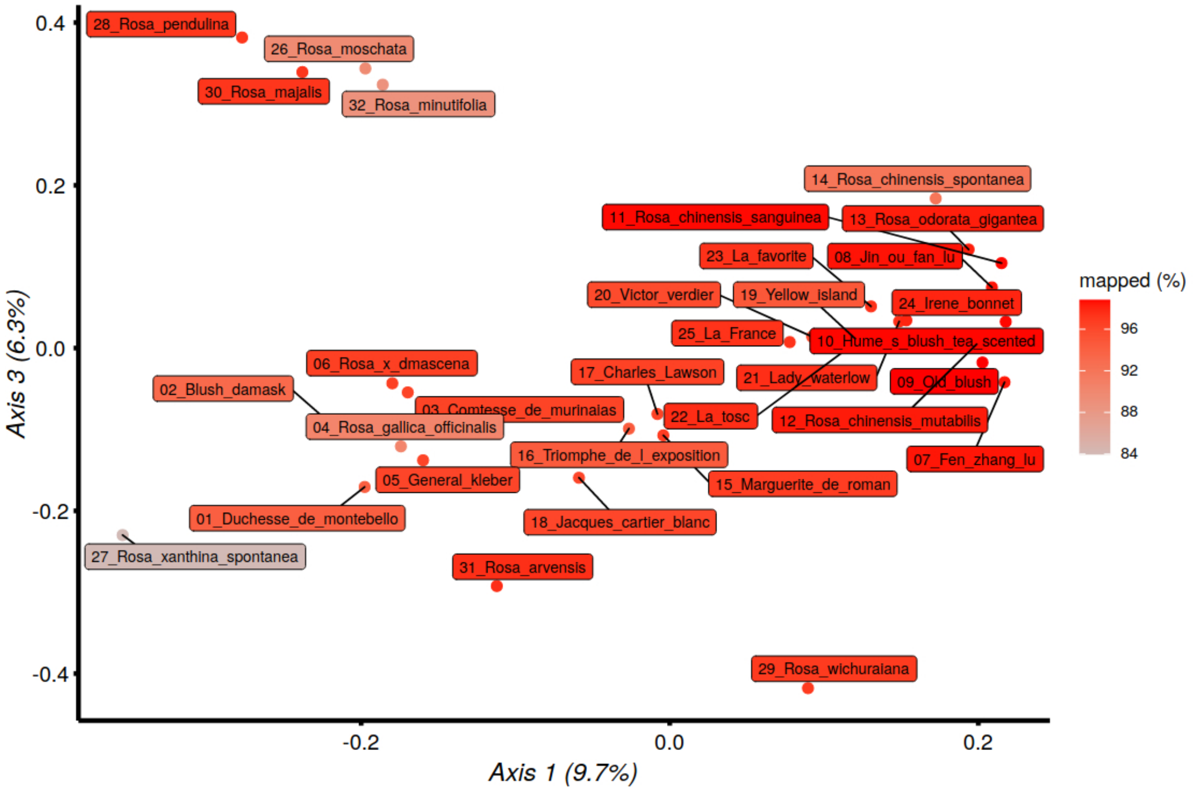

*Figure S3: Mapping rates (proportion of the reads mapped on the 'Old Blush' reference genome, Hibrand-Saint Oyant et al., 2018) for the different samples used in the study, with regards to the observed population structure in Fig. 2B. Despite the use of an ancient Asian reference, the mapping rate is not strongly associated with the two PCs suggesting a limited impact of mapping biases.*

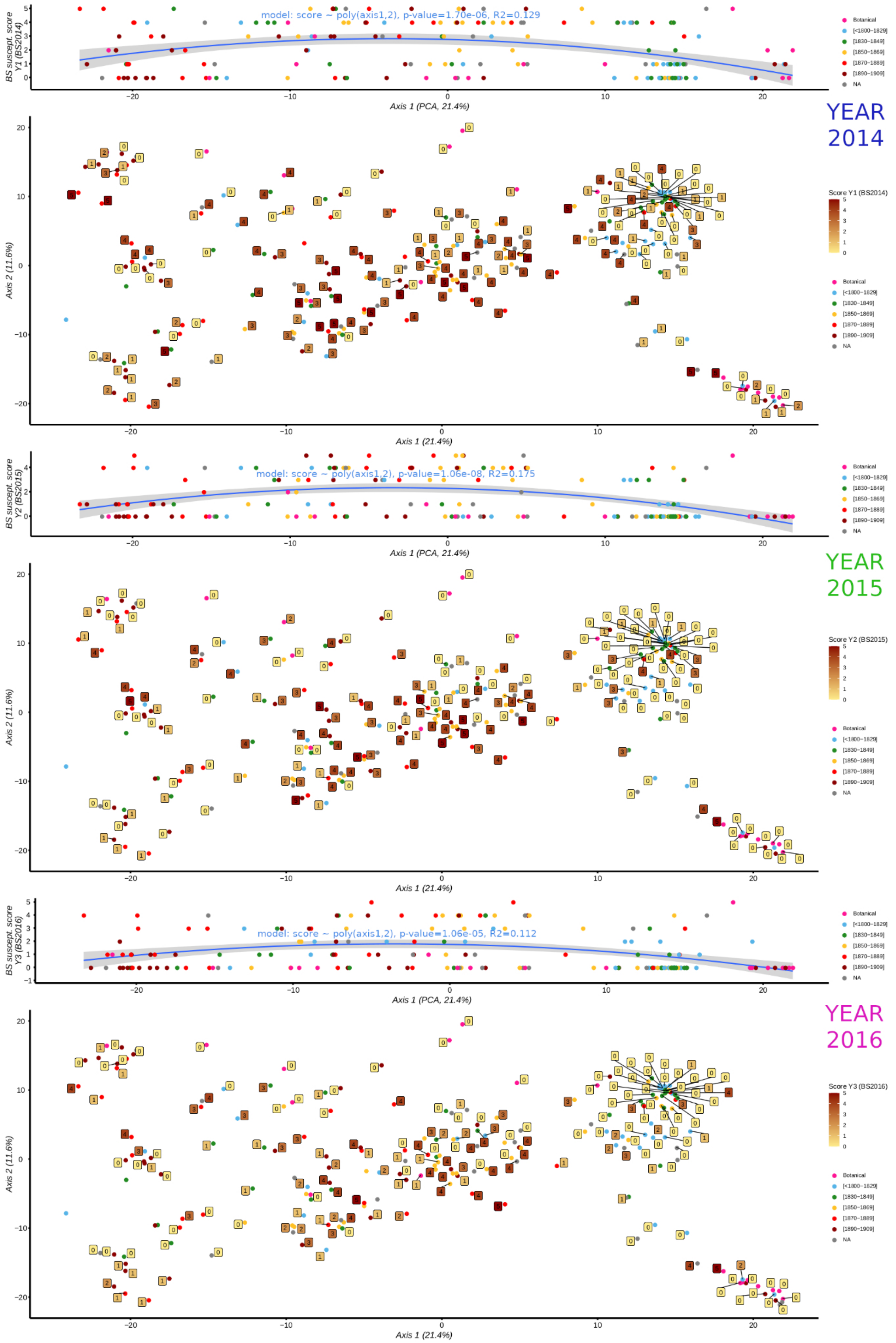
model: score ~ poly(axis1,2), p-value=1.70e-06, R2=0.129
BS suscept. score Y1 (BS2014)
Axis 1 (PCA, 21.4%)
YEAR 2014
Axis 2 (11.6%)
Axis 1 (21.4%)
Score Y1 (BS2014)
Botanical
[<1800–1829]
[1830–1849]
[1850–1869]
[1870–1889]
[1890–1909]
NA
model: score ~ poly(axis1,2), p-value=1.06e-08, R2=0.175
BS suscept. score Y2 (BS2015)
YEAR 2015
Score Y2 (BS2015)
model: score ~ poly(axis1,2), p-value=1.06e-05, R2=0.112
BS suscept. score Y3 (BS2016)
YEAR 2016
Score Y3 (BS2016)

*(continued)*

*Figure S4: Variation of the susceptibility to blackspot disease depending on Asian-European gradient of genetic structure. A, C, E: scatterplots of the blackspot susceptibility score depending on the location of the individual on the PC1 for year 2014, 2015 and 2016, respectively. For each year, a degree-2 polynomial fit is shown and is highly significant (p-value: < 2e-5). B, D, F: two first axes of PCA (as shown in Fig. 2A), but also indicating the susceptibility score as phenotyped each year in the Loubert rose collection. Remarkably, ancient Asian and ancient European exhibit a good level of resistance, while the roses with hybrids between these two groups generally exhibit higher susceptibility scores.*

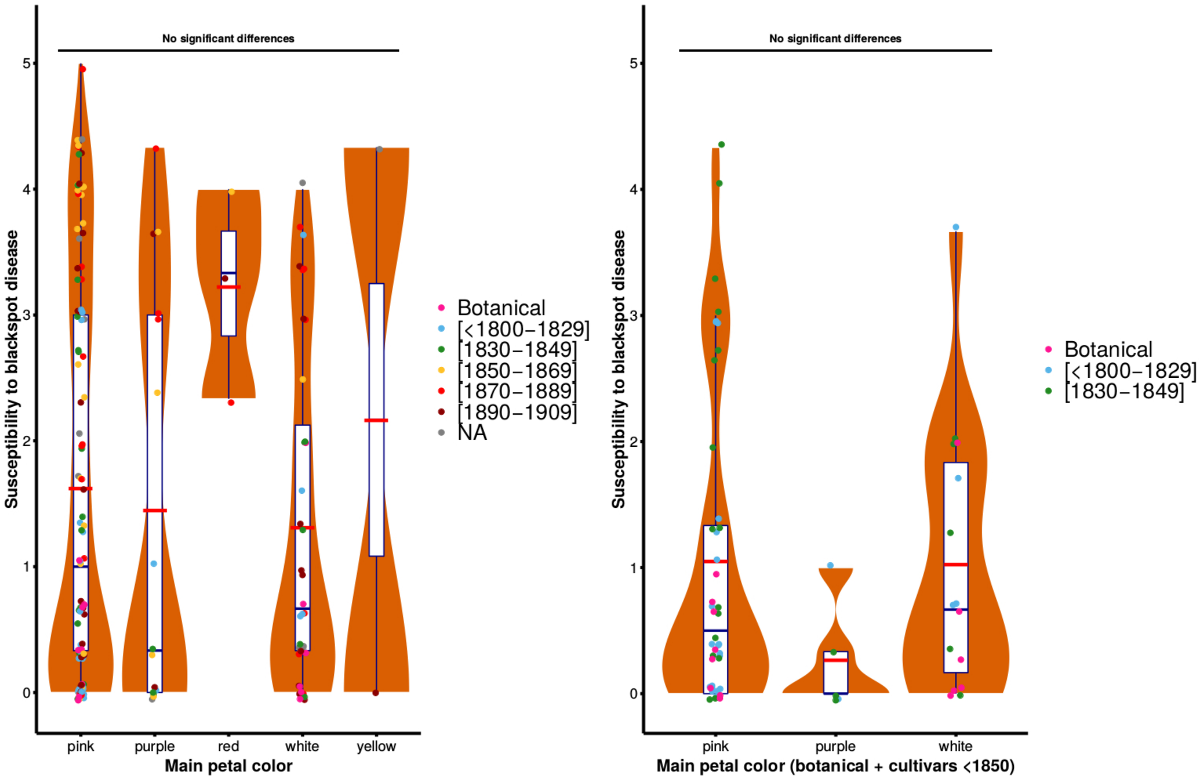

*Figure S5: Black spot susceptibility scores related to the main color of the petals for* ***A)*** *samples having a flower exhibiting a main color (e.g. bicolor flowers were excluded) and* ***B)*** *a subset of samples which corresponds either to botanical accessions or varieties registered before 1850.*

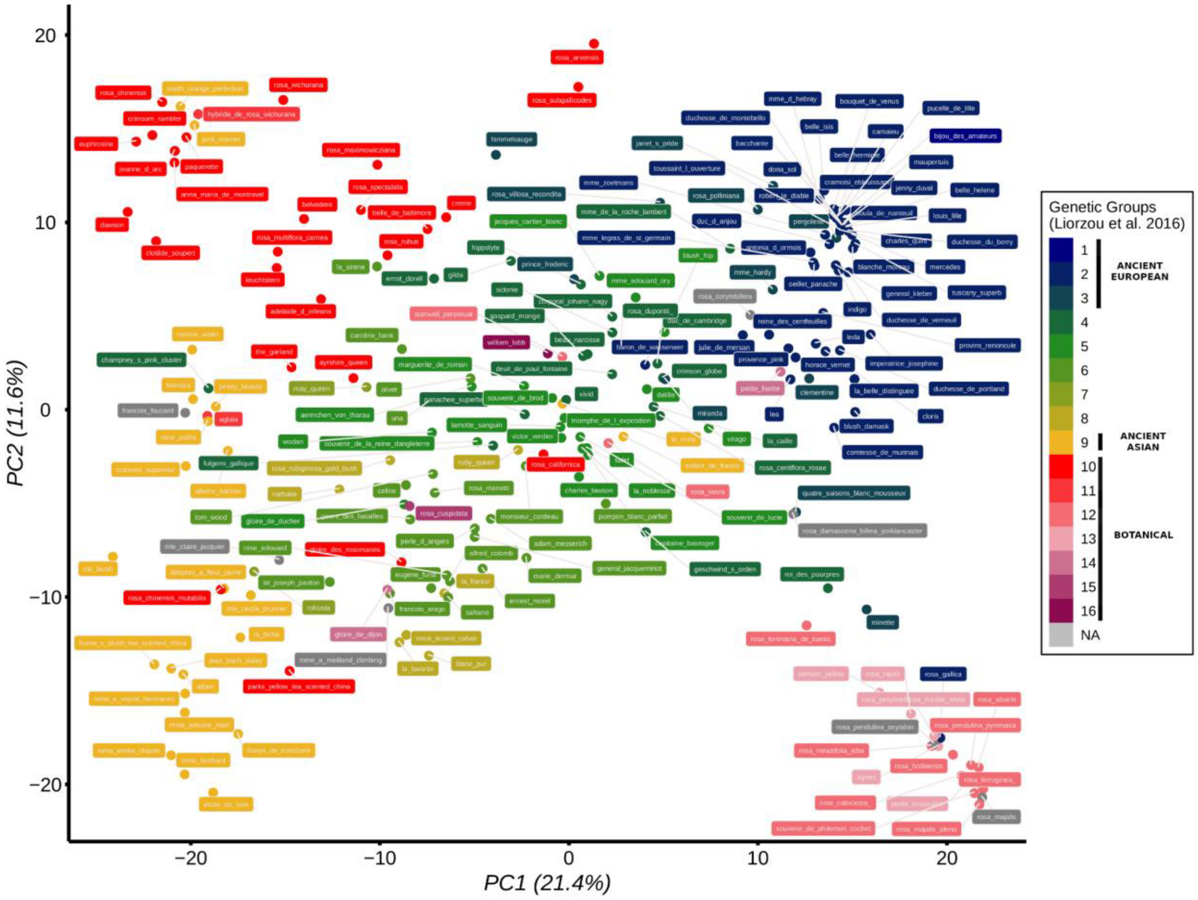

*Figure S6: Population structure of the 204 samples of the SNP array dataset after the exclusion of clones, as shown in Fig. 2A but also some passport data (sample ID, genetic group in Liorzou et al., 2016). The blue and yellow colors correspond to the European and Asian genetic backgrounds, respectively.*

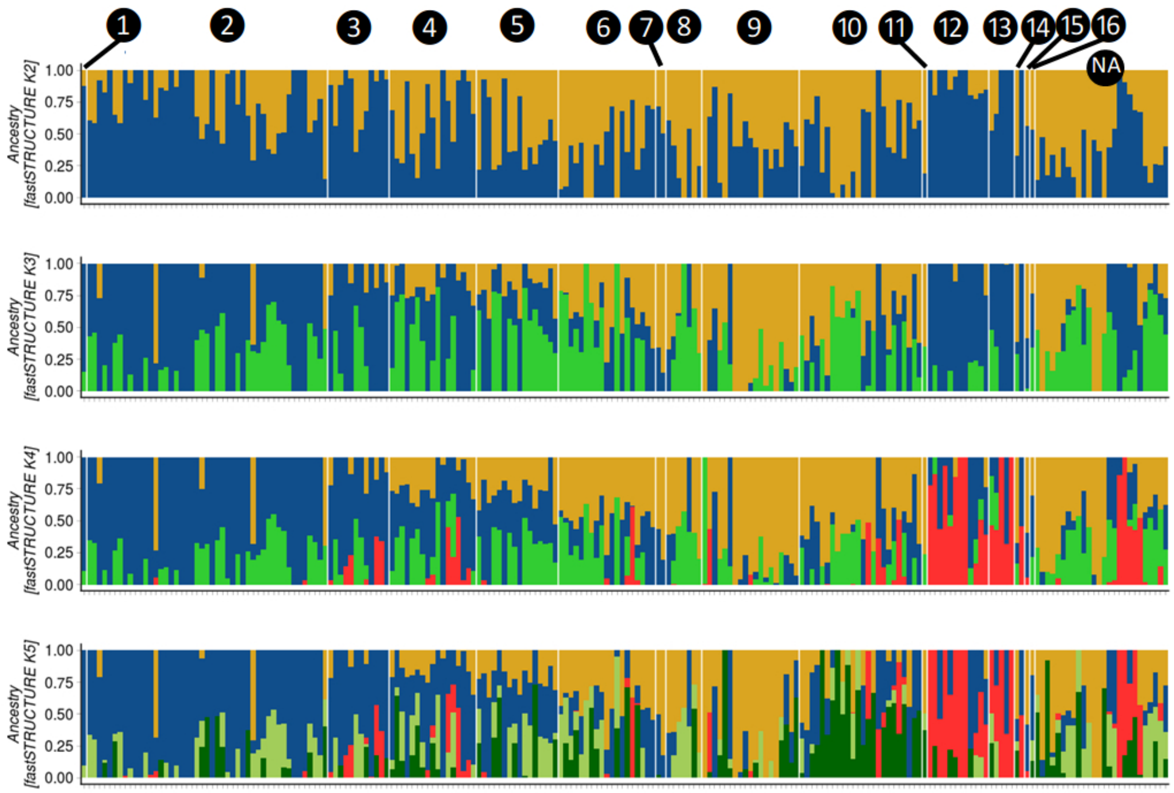

*Figure S7: Ancestry proportions inferred by fastStructure from K=2 to K=5 based on the SNP array data. Two main groups are detected and are consistent with the Asian-European gradient (in yellow and blue, respectively). To check the consistency with a previous work, the membership of the samples to one of the 16 genetic groups as defined in Liorzou et al. (2016) is indicated (except for accessions not previously included).*

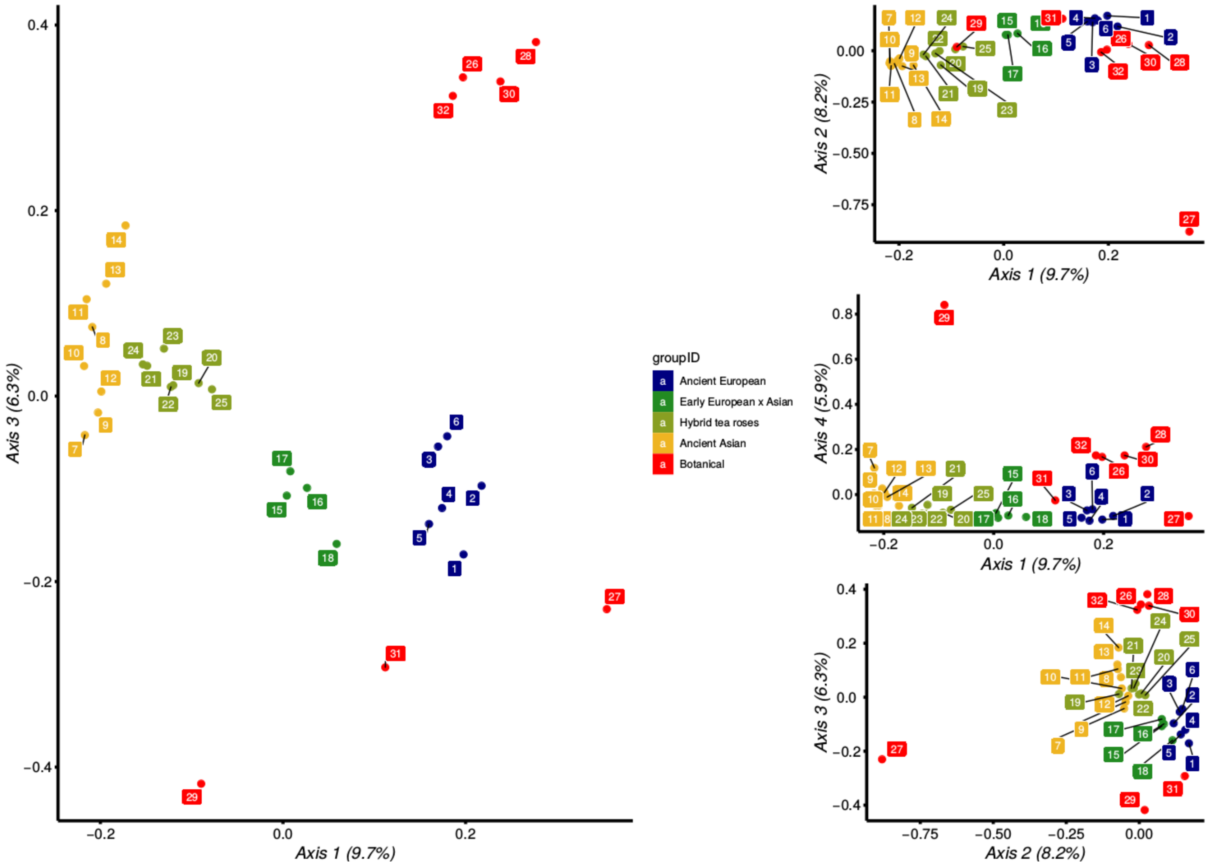

*Figure S8: Four first principal components of a PCA on the 32 whole-genome samples. The main plot corresponds to the one shown in Figure 2B. Individual IDs contain the unique number used in Fig. 2B&C.*

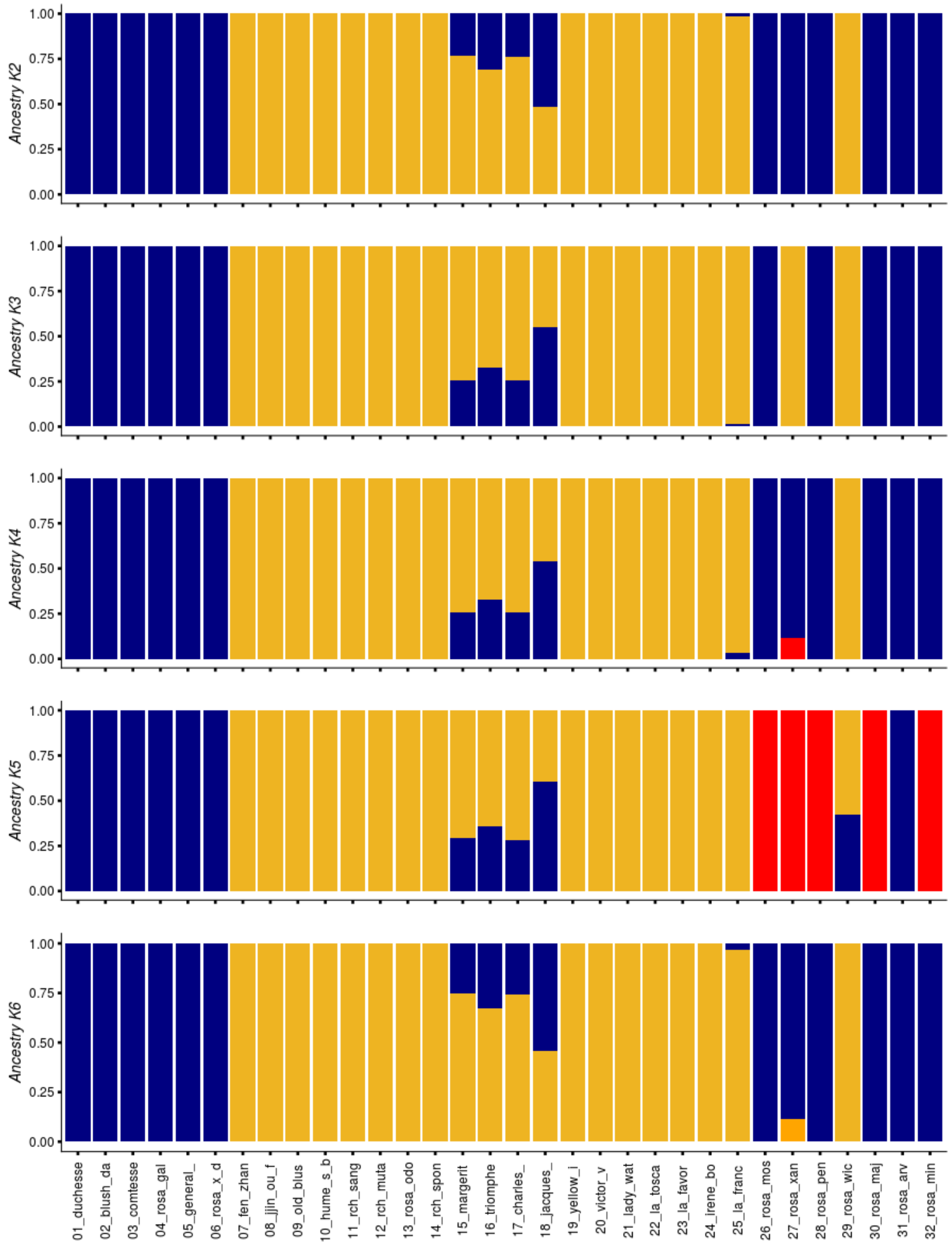

*Figure S9: Ancestry proportions inferred by fastStructure from K=2 to K=6 based on the 50k biallelic SNP derived from WGS data. Two main groups are detected and are consistent with the Asian-European gradient (ancient European: 01-06, ancient Asian: 07-14, early European x Asian: 15-18, Hybrid Tea: 19-35, for more details regarding the accessions, see Table S2). All other groups have extremely low memberships for all individuals ($<1e^{-5}$), explaining why only two or three colors are visible.*

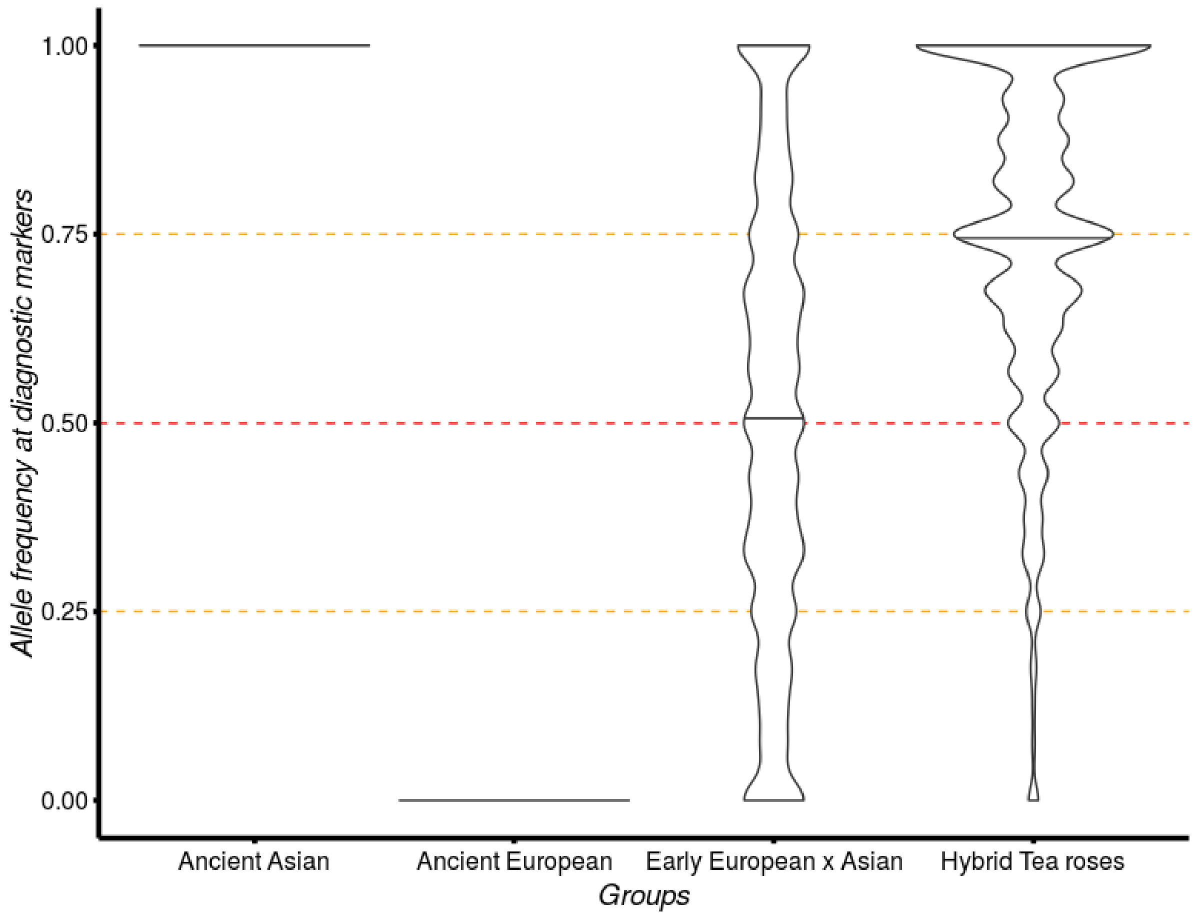

*Figure S10: Distribution of allele frequency at diagnostic loci between our ancient Asian and European samples. The median lines are shown and are here informative about the relative composition of the early European x Asian and Hybrid tea groups with regards to the Asian and European genetic backgrounds. Dotted lines are indicative of allele frequency of 0.25, 0.50 and 0.75. Despite the large variance in the estimate which is inherent to the limited number of individuals used in the study, median values are remarkably close to the 0.50 and 0.75 thresholds for the early European x Asian and the Hybrid tea, respectively. Note that given that the reference rose genome is one of the ancient Asian samples used, diagnostic alleles can only have an allele frequency of 1 and 0 for the reference allele for the ancient Asian and ancient European samples, respectively.*

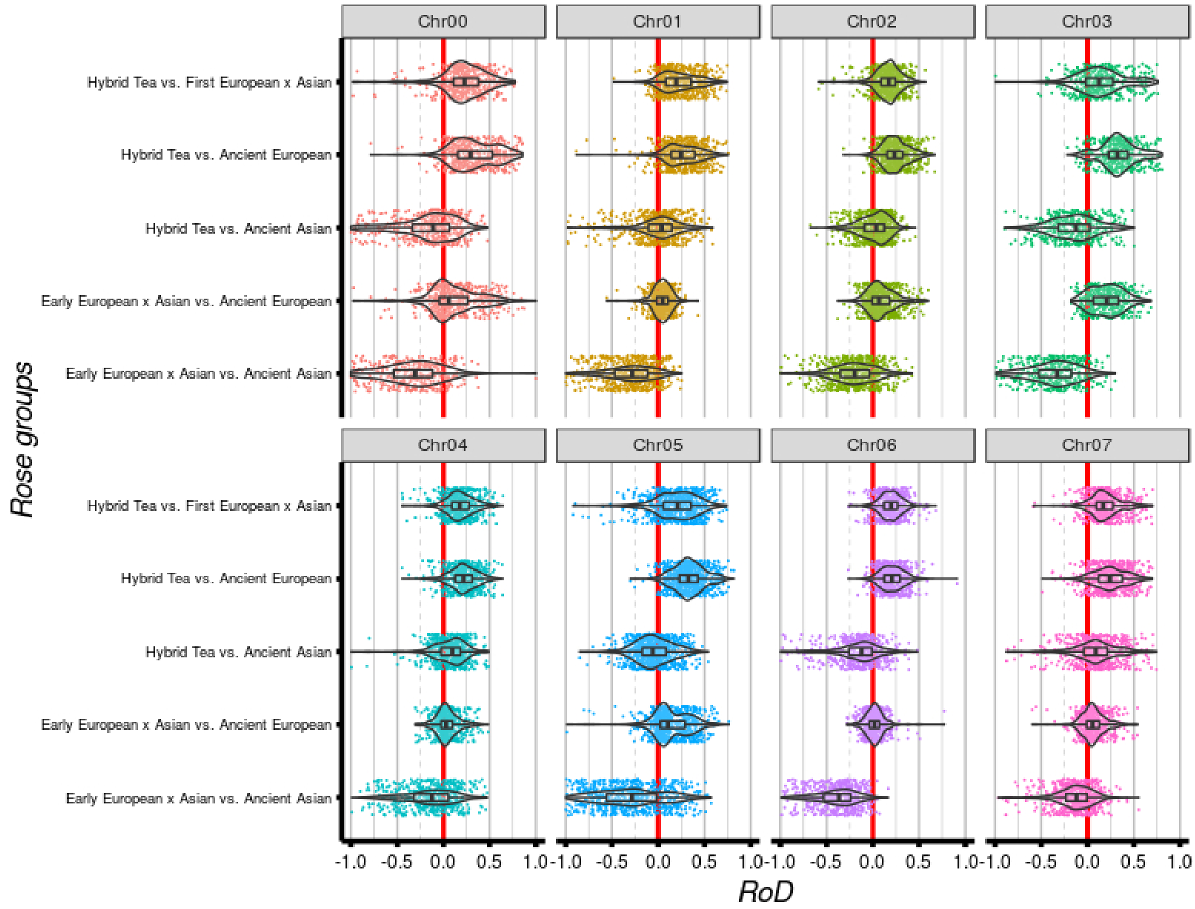

*Figure S11: Distributions of the reduction of diversity (RoD) index per 100 kb windows among groups and chromosomes (from chr01 to chr07, chr 0 corresponds to unanchored sequences). RoD is estimated by comparing the diversity of two groups following the time-series of breeding, with 1 - (π group before / π group after in time). RoD values lower than -1 are not shown. Red line indicates RoD=0 (i.e. same genetic diversity).*

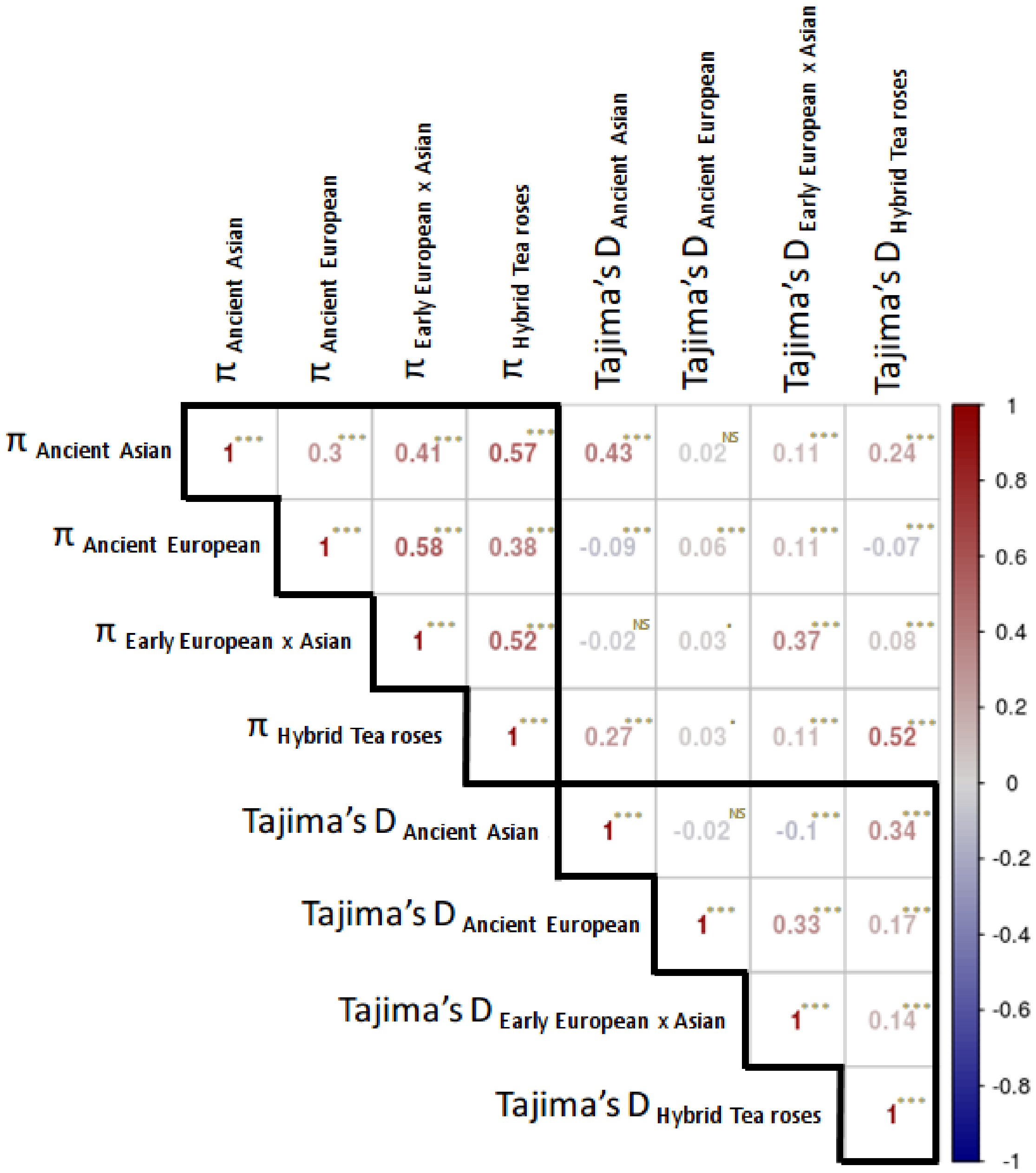

*Figure S12: Pearson's correlation matrix of the nucleotide diversity and Tajima's D genomic landscapes. The color scale ranges from -1 to 1 from blue to red as shown. Significance level: *** < 0.0001, ** < 0.01, * <0.05, · < 0.1, NS non-significant.*

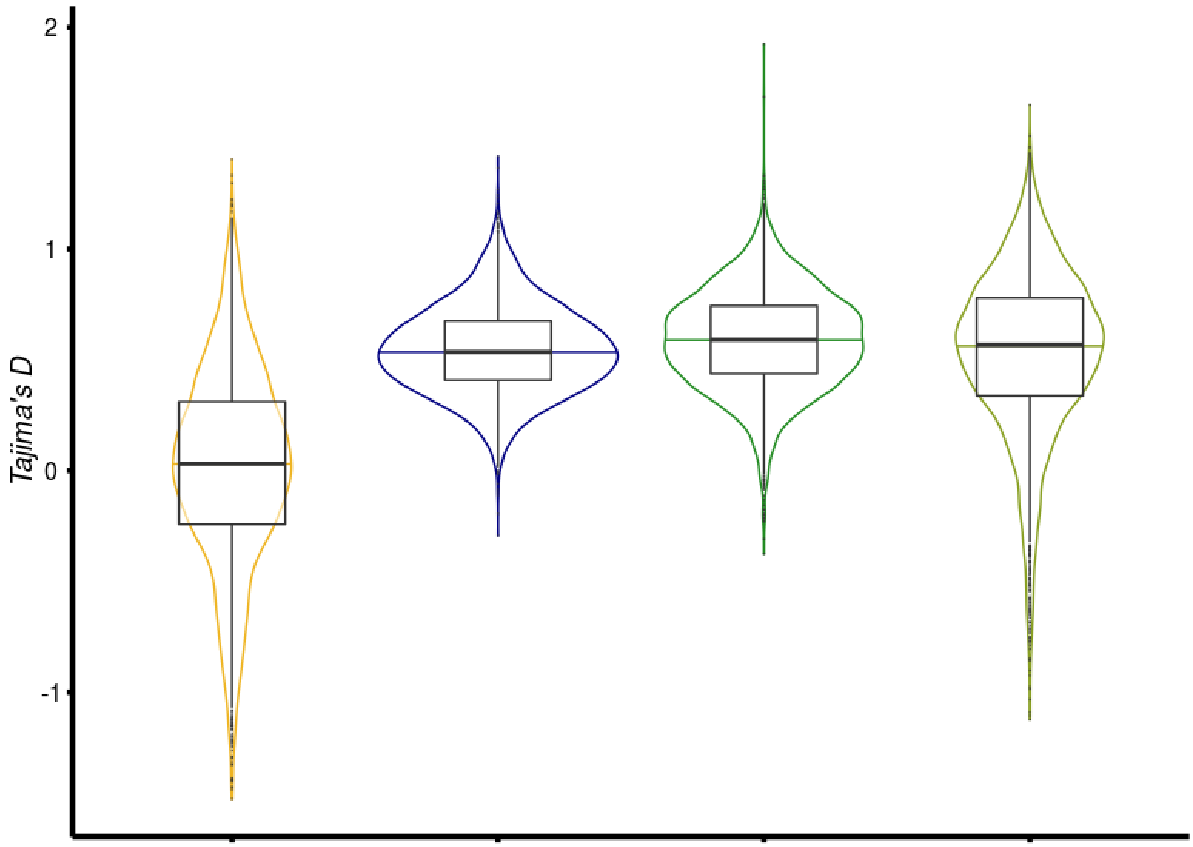

*Figure S13: Distributions of the Tajima's D estimates across the four focal groups based on all 100-kb sliding windows spanning the genome. Ancient Asian, ancient European, early Asian x European and hybrid tea roses are shown in yellow, blue, dark green and khaki, respectively.*

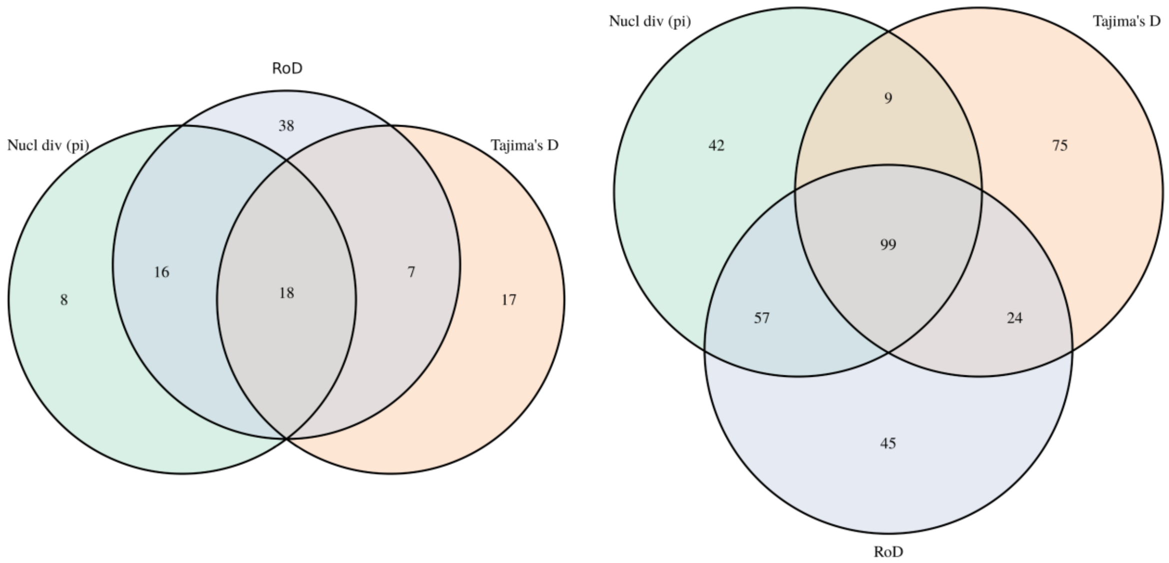

*Fig. S14: Venn diagrams for the impacts of nucleotide diversity (pi, green), Reduction of Diversity (RoD, blue) or Tajima's D (orange) in the detection of regions exhibiting footprints of selection based on* last centile (left) or the five last centiles (right) of the metrics.

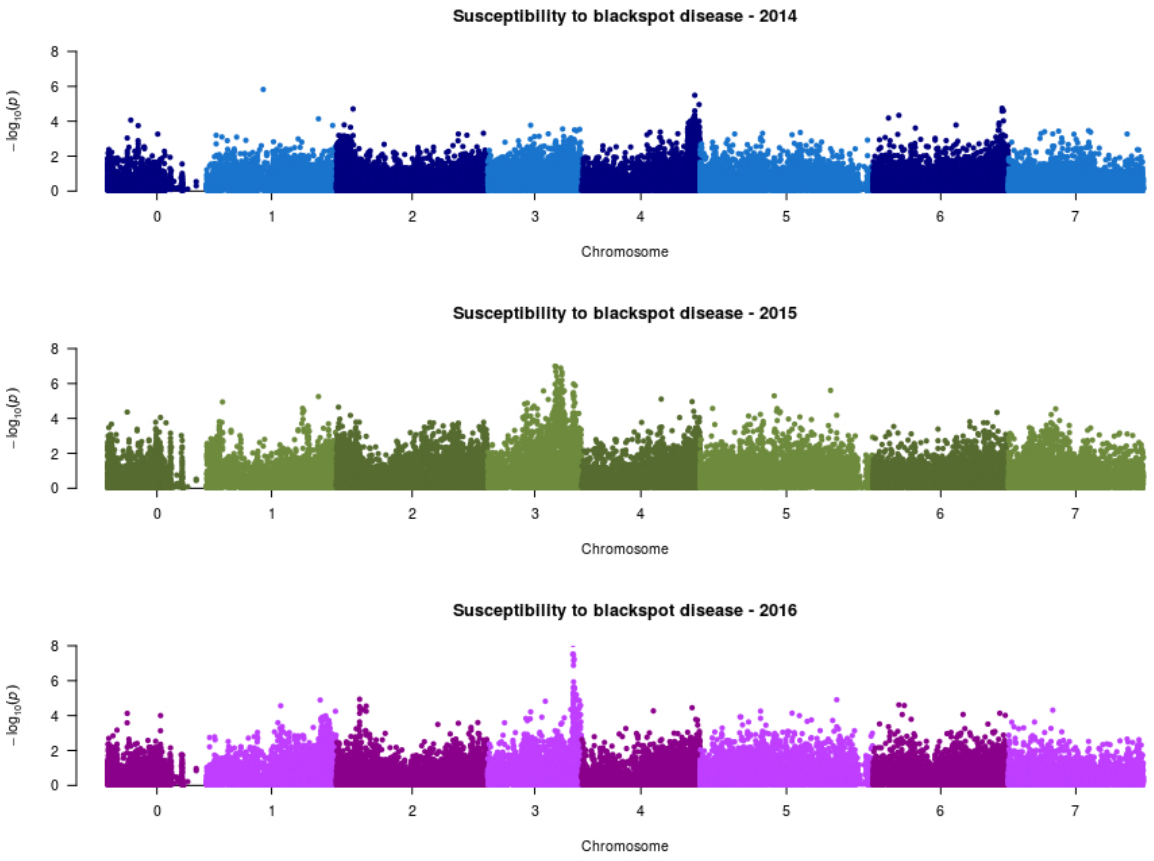

*Fig. S15: Manhattan plots of the -log10(p-values) for the genome-wide associations to susceptibility to blackspot disease in scored years (2014, 2015 and 2016) for the general model of GWASpoly, see also Fig. 4.*

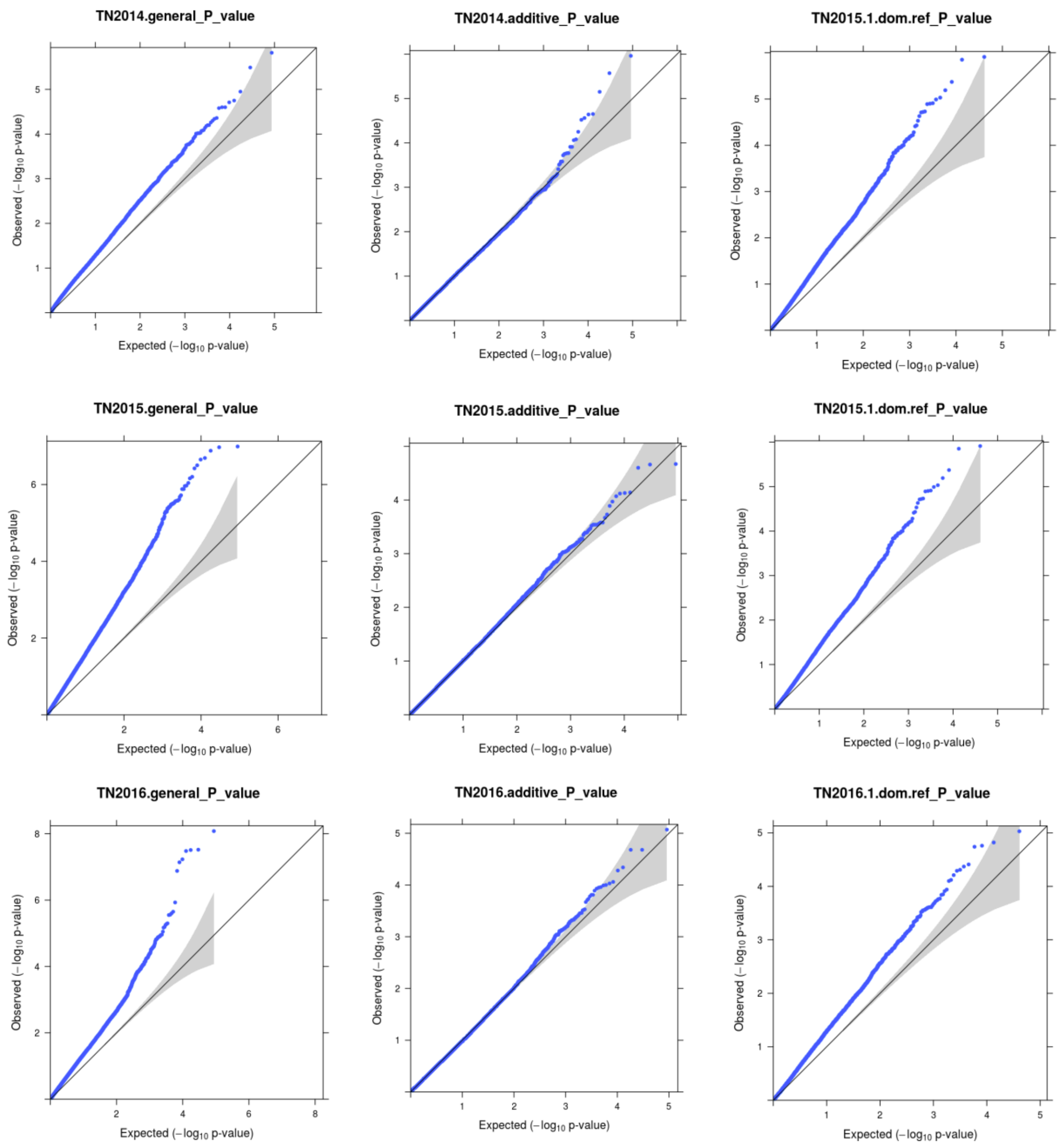

*Fig. S16: QQplots for the GWAS for the scoring of the black spot disease (3 years: 2014 (top), 2015 (middle), 2016 (bottom)), for 3 GWASpoly models (columns), with general (left, as shown in Fig. 4), additive (center) and single dominant for the reference allele (right). Depending on the models, no (e.g. additive models) to substantial inflation of p-values can be observed (e.g. general). All the QQplots are available on the Zenodo repository.*

*Table S1: Potential clones in the dataset, detected through our analyses. The name corresponds to the sample ID, plus the location of the samples in the 96-well plates. Technical duplicate and triplicate samples are shown in yellow and orange, respectively. Other identified clones (unsuspected at the start of the project) are shown in white. Note that all do not necessarily correspond to true clones (consider these genotypes in the other ancient rose gardens), since some could just correspond to mislabelled samples in our local collection.*

| Sample ID1 | Sample ID2 | Sample ID3 (if any) | Sample ID4 (if any) | Sample ID5 (if any) | Sample ID6 (if any) |
|---|---|---|---|---|---|
| quatre_saisons_blanc_mousseux_H6 | quatre_saisons_blanc_mousseux_2_F3 | quatre_saisons_continues_chene_F11 | quatre_saisons_continues_F9 | ispahan_H9 | kazanlik_D5 |
| decoration_de_geschwind_B9 | Decoration_D5 | gilda_A6 | gilda_C10 | geschwind_s_nordland_no_1_H9 | |
| comtesse_de_murinais_E10 | comtesse_de_murinais_2_B1 | zoe_E5 | zoe_2_G4 | berengere_B9 | |
| chloris_B3 | chloris_F10 | felicite_parmentier_C5 | felicite_parmentier_G6 | | |
| adam_2_F4 | adam_A1 | adam_H4 | | | |
| anna_maria_de_montravel_2_F2 | anna_maria_de_montravel_C9 | flocon_de_neige_C4 | | | |
| francois_arago_C6 | francois_arago_2_F11 | prince_noir_F7 | | | |
| belvedere_G8 | princesse_louise_E2 | princesse_marie_B2 | | | |
| celine_2_F12 | celine_A10 | | | | |
| cramoisi_eblouissant_B6 | cramoisi_eblouissant_B5 | | | | |
| ernest_morel_C3 | ernest_morel_2_D4 | | | | |
| ernst_dorell_E8 | ernst_dorell_2_F11 | | | | |
| fatinitza_G9 | fatinitza_2_A3 | | | | |
| horace_vernet_D2 | horace_vernet.E7 | | | | |
| jeanne_d_arc_G2 | jeanne_d_arc_2_G7 | | | | |
| mme_hardy_B6 | mme_hardy_2_B11 | | | | |
| oeillet_panache_G11 | oeillet_panache_2_H12 | | | | |
| persian_yellow_F5 | persian_yellow_A1 | | | | |
| petite_lisette_E3 | petite_lisette_2_E4 | | | | |
| rosa_polliniana_A9 | rosa_polliniana_2_A11 | | | | |
| rosa_villosa_recondita_D2 | rosa_villosa_recondita_2_C5 | | | | |
| amelia_A4 | clementine_B4 | | | | |
| belle_herminie_B10 | kean_D12 | | | | |
| bennet_seedling_A6 | the_garland_D6 | | | | |

| | |
|---|---|
| *bicolore_incomparable_E9* | *crimson_globe_B3* |
| *commandant_beaurepaire_B5* | *panache_superbe_F3* |
| *duchesse_de_montebello_H7* | *narcisse_de_salvandy_E12* |
| *himmelsauge_2_A12* | *rosa_heterophylla_2_A7* |
| *hovyn_de_tronchere_D3* | *mme_falcot_chene_E6* |
| *jean_bach_sisley_D4* | *mme_abel_chatenay_E2* |
| *jenny_duval_G8* | *violacee_E1* |
| *josephine_ritter_C6* | *virago_C8* |
| *la_belle_sultane_D6* | *mercedes_B1* |
| *malvina_D10* | *provence_pink_C2* |
| *mme_edouard_ory_D9* | *mme_louis_leveque_E10* |
| *mme_lierval_E9* | *sir_joseph_paxton_C1* |
| *mme_louise_odier_E11* | *perle_d_angers_F12* |
| *mogador_F2* | *stanwell_perpetual_D11* |
| *rosa_pimpinellifolia_double_white_F6* | *staffa_E11* |
| *ruby_queen_H3* | *triomphe_de_caen_chene_G11* |

*Table S2: Accessions used in this study for the WGS dataset. The accession number shown in the first column corresponds to the IDs used in Figs. 2B and 2C. For more details regarding the results that have led to the inference of the ploidy level, see Extended Fig. S2 (also available "*Extended_FigS2_ploidy_chr_per_chr.pdf*" in the Zenodo repository). For some individuals, this inference is complex, either because of the low coverage or the complex patterns (e.g. some botanical roses).*

| **Acc. Number** | **Accession** | **Group** | **Sample_ID (short)** | **Seq. runs (SRA)** | **Publication** | **Included for nucl. diversity (reason)** | **Inferred ploidy** | **Mean cov (at SNP pos)** |
|---|---|---|---|---|---|---|---|---|
| 1 | Duchesse de montebello | Ancient European | 175580 | SRR25401352 | This study | No (1st level kinship) | 4 | 10.63 |
| 2 | Blush damask | Ancient European | 175581 | SRR25401353 | This study | **Yes** | 4 | 8.88 |
| 3 | Comtesse de murinais | Ancient European | 175583 | SRR25401344 | This study | **Yes** | 4 | 8.18 |
| 4 | Rosa gallica officinalis | Ancient European | Rgaloff | SRR7077016 | Hibrand-Saint Oyant *et al.*, 2018 | **Yes** | 4 | 44.47 |
| 5 | General kleber | Ancient European | 177142 | SRR25401341 | This study | **Yes** | 4 | 6.72 |
| 6 | Rosa x damascena | Ancient European | Rxdama | SRR6175509, SRR6175510 | Raymond *et al.*, 2018 | No (1st level kinship) | 4 | 15.72 |

| | | | | | | | | |
|---|---|---|---|---|---|---|---|---|
| 7 | Fen zhang lu | Ancient Asian | 175584 | SRR25401345 | This study | **Yes** | 2 | 13.95 |
| 8 | Jin ou fan lu | Ancient Asian | 175585 | SRR25401346 | This study | **Yes** | 4 | 9.01 |
| 9 | Old blush | Ancient Asian | OB2n | SRR25401343 | This study | **Yes** | 2 | 43.50 |
| 10 | Hume's blush tea scented | Ancient Asian | HumeBlush | SRR6175507 | Raymond *et al.*, 2018 | No (1st level kinship) | 2 | 20.29 |
| 11 | Rosa chinensis sanguinea | Ancient Asian | Rchisan | SRR6175515 | Raymond *et al.*, 2018 | **Yes** | 2 | 19.67 |
| 12 | Rosa chinensis mutabilis | Ancient Asian | Rchimut | SRR6175518 | Raymond *et al.*, 2018 | **Yes** | 2 | 19.00 |
| 13 | Rosa odorata gigantea | Ancient Asian | Rogig | SRR6175516, SRR6175517 | Raymond *et al.*, 2018 | **Yes** | 2 | 16.74 |
| 14 | Rosa chinensis spontanea | Ancient Asian | Rchispo | SRR7077020 | Hibrand-Saint Oyant *et al.*, 2018 | **Yes** | 2 | 30.96 |

| | | | | | | | | |
|---|---|---|---|---|---|---|---|---|
| 15 | Margerite_de_roman | Early European x Asian | 175586 | SRR25401347 | This study | **Yes** | 4 | 6.83 |
| 16 | Triomphe de l'exposition | Early European x Asian | 175587 | SRR25401348 | This study | **Yes** | 4 | 7.90 |
| 17 | Charles Lawson | Early European x Asian | 175623 | SRR25401350 | This study | **Yes** | 4 | 7.19 |
| 18 | Jacques Cartier blanc | Early European x Asian | 175624 | SRR25401351 | This study | **Yes** | 4 | 6.77 |
| 19 | Yellow island | Hybrid tea roses | YellowIsland | SRR10037956, SRR10037957, SRR10037958 | Unpublished | No (too modern variety) | 4 | 33.92 |
| 20 | Victor Verdier | Hybrid tea roses | 175588 | SRR25401349 | This study | No (uncertainty*) | 4 | 5.19 |
| 21 | Lady Waterlow | Hybrid tea roses | 175626 | SRR25401354 | This study | **Yes** | 4 | 6.99 |
| 22 | La tosca | Hybrid tea roses | 175627 | SRR25401339 | This study | **Yes** | 4 | 8.14 |

| | | | | | | | | |
|---|---|---|---|---|---|---|---|---|
| 23 | La favorite | Hybrid tea roses | 175628 | SRR25401340 | This study | **Yes** | 4 | 7.10 |
| 24 | Irene Bonnet | Hybrid tea roses | 177143 | SRR25401342 | This study | **Yes** | 4 | 10.17 |
| 25 | La France | Hybrid tea roses | LaFrance | SRR6175521 | Raymond *et al.*, 2018 | No (triploidy) | 3 | 29.15 |
| 26 | Rosa moschata | Botanical | Rmosch | SRR7077017 | Hibrand-Saint Oyant *et al.*, 2018 | No (botanical) | 2 | 20.99 |
| 27 | Rosa xanthina spontanea | Botanical | Rxanth | SRR7077022 | Hibrand-Saint Oyant *et al.*, 2018 | No (botanical) | 2 | 19.65 |
| 28 | Rosa pendulina | Botanical | Rpend | SRR6175522 | Raymond *et al.*, 2018 | No (botanical) | 4 | 27.26 |
| 29 | Rosa wichuraiana | Botanical | Rwich | SRR6175519 & SRR6175520 | Raymond *et al.*, 2018 | No (botanical) | 2 | 17.20 |
| 30 | Rosa majalis | Botanical | Rmaja | SRR6175513 | Raymond *et al.*, 2018 | No (botanical) | 4 | 19.61 |

| | | | | | | | | |
|---|---|---|---|---|---|---|---|---|
| 31 | Rosa arvensis | Botanical | Rarve | SRR6175512 | Raymond *et al.*, 2018 | No (botanical) | 4 | 18.70 |
| 32 | Rosa minutifolia | Botanical | Rminut | SRR7077023 | Hibrand-Saint Oyant *et al.*, 2018 | No (botanical) | 2 | 20.81 |

*Table S3: number of diagnostic alleles per chromosome. Last column indicates the average number of diagnostic SNPs per Mb on the corresponding chromosomes.*

| | # Diagnostic SNPs | Chromosome length | # Diagnostic/Mb |
|---|---|---|---|
| Chr00 | 10,558 | 52,404,850 | 201.6 |
| Chr01 | 15,487 | 64,770,848 | 239.1 |
| Chr02 | 15,873 | 75,129,302 | 211.3 |
| Chr03 | 30,235 | 46,843,630 | 645.4 |
| Chr04 | 17,585 | 59,004,735 | 298.0 |
| Chr05 | 32,184 | 85,885,663 | 374.7 |
| Chr06 | 23,785 | 67,395,200 | 352.9 |
| Chr07 | 24,930 | 67,081,725 | 371.6 |
| Total | 170,637 | 518,515,953 | 329.1 |

*Table S4: List of 100 kb windows exhibiting* a shared signal of low π, negative Tajima's D and high RoD when $\pi_{\text{hybrid tea roses}}$ is compared with either $\pi_{\text{Asian}}$ or $\pi_{\text{European}}$. All windows are in the top 5% criteria used to detect *footprints of artificial selection (see main text), but windows associated with the most stringent filtering criteria (top 1%) are shown in bold. Differences in Tajima's D between Hybrid Tea and Ancient Asian samples (Asia) or ancient European samples (Europe) correspond to absolute values.*

| ***Chr.*** | ***startwind.*** | ***end wind.*** | ***π Asia*** | ***Tajima's*** $D_{Asia}$ | ***π Europe*** | ***Tajima*** $D_{Europe}$ | ***π HybridTea*** | $D_{HybridTea}$ | ***RoD Hybrid Tea vs. Asia*** | ***RoD Hybrid Tea vs. Europe*** | ***ΔTajima's D HybridTea vs. Asia*** | ***ΔTajima's D HybridTea vs. Europe*** |
|---|---|---|---|---|---|---|---|---|---|---|---|---|
| *Chr01* | *32.1* | *32.2* | *1.13E-02* | *-0.1701* | *1.67E-02* | *0.7973* | *5.43E-03* | *-0.3619* | *0.5200* | *0.6755* | *0.1918* | *1.1592* |
| *Chr01* | *32.4* | *32.5* | *1.24E-02* | *-0.5144* | *1.50E-02* | *0.4267* | *5.86E-03* | *-0.4724* | *0.5277* | *0.6100* | *0.0420* | *0.8991* |
| *Chr01* | *32.5* | *32.6* | *9.77E-03* | *-0.4538* | *2.15E-02* | *0.7461* | *5.23E-03* | *-0.1927* | *0.4646* | *0.7572* | *0.2611* | *0.9388* |
| *Chr01* | *32.6* | *32.7* | *8.49E-03* | *-0.6235* | *1.55E-02* | *0.3452* | *4.95E-03* | *-0.5166* | *0.4169* | *0.6803* | *0.1069* | *0.8618* |
| *Chr01* | *33.5* | *33.6* | *1.14E-02* | *-0.3170* | *1.66E-02* | *0.5249* | *6.47E-03* | *-0.6335* | *0.4310* | *0.6111* | *0.3165* | *1.1584* |
| ***Chr01*** | ***33.6*** | ***33.7*** | ***1.09E-02*** | ***-0.5408*** | ***1.72E-02*** | ***0.8522*** | ***4.49E-03*** | ***-0.7713*** | ***0.5898*** | ***0.7383*** | ***0.2305*** | ***1.6235*** |
| *Chr01* | *34.3* | *34.4* | *8.28E-03* | *-0.2913* | *1.58E-02* | *0.4375* | *5.08E-03* | *-0.5987* | *0.3870* | *0.6780* | *0.3075* | *1.0362* |
| *Chr01* | *34.6* | *34.7* | *8.18E-03* | *-0.2413* | *1.42E-02* | *0.4708* | *4.87E-03* | *-0.3361* | *0.4048* | *0.6577* | *0.0948* | *0.8068* |
| *Chr01* | *36.3* | *36.4* | *7.48E-03* | *-0.2690* | *1.92E-02* | *0.7908* | *5.71E-03* | *-0.4890* | *0.2376* | *0.7035* | *0.2200* | *1.2798* |
| *Chr01* | *36.4* | *36.5* | *1.31E-02* | *-0.1596* | *1.36E-02* | *0.7253* | *5.54E-03* | *-0.8506* | *0.5771* | *0.5920* | *0.6911* | *1.5759* |
| *Chr01* | *39.5* | *39.6* | *1.23E-02* | *-0.0444* | *1.63E-02* | *0.5327* | *6.88E-03* | *-0.6205* | *0.4424* | *0.5787* | *0.5761* | *1.1532* |
| *Chr01* | *39.9* | *40* | *1.61E-02* | *-0.1864* | *1.90E-02* | *0.8274* | *7.25E-03* | *-0.7581* | *0.5491* | *0.6189* | *0.5717* | *1.5855* |
| *Chr01* | *40.1* | *40.2* | *1.12E-02* | *-0.2852* | *1.84E-02* | *0.4577* | *6.72E-03* | *-0.5840* | *0.3980* | *0.6359* | *0.2987* | *1.0417* |
| *Chr01* | *40.8* | *40.9* | *1.31E-02* | *-0.3184* | *1.88E-02* | *0.3943* | *6.78E-03* | *-0.6205* | *0.4812* | *0.6399* | *0.3022* | *1.0149* |
| *Chr01* | *42.4* | *42.5* | *9.53E-03* | *-0.3140* | *1.96E-02* | *0.7245* | *7.11E-03* | *-0.4644* | *0.2537* | *0.6381* | *0.1504* | *1.1889* |
| *Chr01* | *43.1* | *43.2* | *7.91E-03* | *-0.6466* | *1.62E-02* | *0.4253* | *6.19E-03* | *-0.3465* | *0.2179* | *0.6183* | *0.3001* | *0.7718* |
| *Chr01* | *63.6* | *63.7* | *7.16E-03* | *-0.6452* | *1.56E-02* | *0.7996* | *4.27E-03* | *-0.4881* | *0.4041* | *0.7266* | *0.1571* | *1.2876* |
| *Chr02* | *27.3* | *27.4* | *1.29E-02* | *-0.0327* | *1.66E-02* | *0.2588* | *6.97E-03* | *-0.2505* | *0.4598* | *0.5795* | *0.2178* | *0.5093* |
| *Chr02* | *36.5* | *36.6* | *4.88E-03* | *-0.6467* | *1.41E-02* | *0.4052* | *6.09E-03* | *-0.2013* | *-0.2474* | *0.5672* | *0.4454* | *0.6065* |
| *Chr02* | *41.1* | *41.2* | *8.51E-03* | *-0.5741* | *1.58E-02* | *0.5774* | *5.42E-03* | *-0.2216* | *0.3637* | *0.6578* | *0.3525* | *0.7990* |
| *Chr03* | *24.4* | *24.5* | *8.63E-03* | *-0.1338* | *1.93E-02* | *0.5796* | *5.98E-03* | *-0.6392* | *0.3073* | *0.6899* | *0.5054* | *1.2187* |
| ***Chr03*** | ***24.6*** | ***24.7*** | ***6.29E-03*** | ***-0.5887*** | ***1.59E-02*** | ***0.5145*** | ***4.75E-03*** | ***-0.6262*** | ***0.2445*** | ***0.7000*** | ***0.0375*** | ***1.1407*** |
| ***Chr03*** | ***25.0*** | ***25.1*** | ***5.86E-03*** | ***-1.0113*** | ***1.70E-02*** | ***0.6488*** | ***4.12E-03*** | ***-0.7520*** | ***0.2971*** | ***0.7577*** | ***0.2593*** | ***1.4008*** |
| ***Chr03*** | ***25.1*** | ***25.2*** | ***5.35E-03*** | ***-1.3947*** | ***1.65E-02*** | ***0.3351*** | ***3.90E-03*** | ***-1.1065*** | ***0.2719*** | ***0.7631*** | ***0.2882*** | ***1.4416*** |

| | | | | | | | | | | | | |
|---|---|---|---|---|---|---|---|---|---|---|---|---|
| **Chr03** | **25.2** | **25.3** | **7.07E-03** | **-1.1096** | **1.65E-02** | **0.2086** | **4.46E-03** | **-0.9759** | **0.3697** | **0.7303** | **0.1337** | **1.1845** |
| **Chr03** | **25.3** | **25.4** | **5.57E-03** | **-1.2992** | **1.67E-02** | **0.6086** | **3.61E-03** | **-1.1236** | **0.3520** | **0.7836** | **0.1755** | **1.7322** |
| **Chr03** | **25.4** | **25.5** | **5.70E-03** | **-1.0114** | **1.53E-02** | **0.5271** | **3.66E-03** | **-1.0329** | **0.3572** | **0.7609** | **0.0215** | **1.5600** |
| **Chr03** | **25.6** | **25.7** | **6.70E-03** | **-1.0002** | **1.69E-02** | **0.4024** | **4.14E-03** | **-0.8562** | **0.3833** | **0.7548** | **0.1439** | **1.2586** |
| **Chr03** | **26.0** | **26.1** | **5.50E-03** | **-0.6393** | **1.72E-02** | **0.4806** | **4.44E-03** | **-0.7167** | **0.1936** | **0.7421** | **0.0774** | **1.1974** |
| Chr03 | 26.1 | 26.2 | 9.04E-03 | -0.1546 | 1.67E-02 | 0.3360 | 5.44E-03 | -0.8477 | 0.3986 | 0.6751 | 0.6931 | 1.1837 |
| Chr03 | 26.2 | 26.3 | 9.09E-03 | -0.1002 | 1.74E-02 | 0.2270 | 5.72E-03 | -0.7661 | 0.3705 | 0.6710 | 0.6659 | 0.9931 |
| **Chr03** | **26.5** | **26.6** | **8.32E-03** | **-0.3158** | **1.84E-02** | **0.5088** | **4.14E-03** | **-0.9249** | **0.5027** | **0.7751** | **0.6091** | **1.4337** |
| **Chr03** | **26.7** | **26.8** | **5.97E-03** | **-0.6281** | **1.52E-02** | **0.2159** | **3.88E-03** | **-0.9849** | **0.3497** | **0.7447** | **0.3568** | **1.2008** |
| Chr03 | 26.8 | 26.9 | 6.01E-03 | -0.3066 | 1.67E-02 | 0.2804 | 5.19E-03 | -0.5402 | 0.1362 | 0.6899 | 0.2336 | 0.8206 |
| Chr03 | 26.9 | 27 | 6.67E-03 | -0.4732 | 1.57E-02 | 0.2930 | 5.39E-03 | -0.4914 | 0.1917 | 0.6558 | 0.0182 | 0.7844 |
| **Chr03** | **27.0** | **27.1** | **6.90E-03** | **-0.8315** | **1.59E-02** | **0.3975** | **3.57E-03** | **-0.8559** | **0.4828** | **0.7757** | **0.0244** | **1.2534** |
| **Chr03** | **27.1** | **27.2** | **5.72E-03** | **-0.9526** | **1.84E-02** | **0.4980** | **3.91E-03** | **-0.9014** | **0.3168** | **0.7881** | **0.0511** | **1.3995** |
| **Chr03** | **27.4** | **27.5** | **5.63E-03** | **-0.3124** | **1.74E-02** | **0.6507** | **4.77E-03** | **-0.8051** | **0.1517** | **0.7262** | **0.4927** | **1.4558** |
| Chr03 | 27.6 | 27.7 | 6.81E-03 | -0.7113 | 1.86E-02 | 0.4422 | 4.47E-03 | -0.6071 | 0.3442 | 0.7604 | 0.1042 | 1.0493 |
| Chr03 | 27.7 | 27.8 | 7.39E-03 | -0.3291 | 1.60E-02 | 0.3509 | 5.28E-03 | -0.5024 | 0.2852 | 0.6710 | 0.1732 | 0.8533 |
| Chr03 | 27.8 | 27.9 | 8.57E-03 | -0.1309 | 1.69E-02 | 0.2692 | 6.28E-03 | -0.3833 | 0.2668 | 0.6275 | 0.2525 | 0.6525 |
| **Chr03** | **28.0** | **28.1** | **4.97E-03** | **-1.1854** | **1.80E-02** | **0.1137** | **3.47E-03** | **-1.0910** | **0.3024** | **0.8076** | **0.0944** | **1.2047** |
| **Chr03** | **28.1** | **28.2** | **6.07E-03** | **-0.6816** | **1.81E-02** | **0.3090** | **4.75E-03** | **-0.7906** | **0.2184** | **0.7374** | **0.1090** | **1.0997** |
| Chr03 | 32.5 | 32.6 | 6.16E-03 | -0.2263 | 1.47E-02 | 0.3759 | 5.11E-03 | -0.3795 | 0.1715 | 0.6532 | 0.1532 | 0.7554 |
| Chr03 | 32.7 | 32.8 | 6.92E-03 | -0.8530 | 1.88E-02 | 0.6716 | 5.60E-03 | -0.4552 | 0.1895 | 0.7016 | 0.3978 | 1.1268 |
| Chr03 | 32.8 | 32.9 | 7.61E-03 | -0.3849 | 1.68E-02 | 0.4918 | 6.76E-03 | -0.2260 | 0.1119 | 0.5986 | 0.1589 | 0.7178 |
| Chr03 | 33.0 | 33.1 | 8.37E-03 | -0.8407 | 1.78E-02 | 0.2977 | 6.19E-03 | -0.5176 | 0.2602 | 0.6513 | 0.3231 | 0.8152 |
| Chr04 | 47.4 | 47.5 | 5.94E-03 | -1.2927 | 1.47E-02 | 0.7838 | 6.00E-03 | -0.3529 | -0.0099 | 0.5909 | 0.9398 | 1.1367 |
| Chr04 | 48.1 | 48.2 | 8.10E-03 | -1.1516 | 1.67E-02 | 0.5244 | 6.04E-03 | -0.3455 | 0.2547 | 0.6388 | 0.8061 | 0.8699 |
| Chr04 | 48.7 | 48.8 | 7.41E-03 | -1.2313 | 1.95E-02 | 0.8907 | 7.13E-03 | -0.3600 | 0.0373 | 0.6340 | 0.8714 | 1.2506 |
| Chr05 | 15.9 | 16 | 9.19E-03 | -0.2760 | 1.63E-02 | 0.2556 | 5.97E-03 | -0.3006 | 0.3500 | 0.6332 | 0.0246 | 0.5562 |
| Chr05 | 16.4 | 16.5 | 9.37E-03 | 0.0770 | 1.44E-02 | 0.5987 | 4.30E-03 | -0.5925 | 0.5414 | 0.7021 | 0.6695 | 1.1912 |
| Chr05 | 16.7 | 16.8 | 9.51E-03 | -0.6503 | 1.64E-02 | 0.3777 | 5.52E-03 | -0.7516 | 0.4201 | 0.6629 | 0.1013 | 1.1293 |

| | | | | | | | | | | | | |
|---|---|---|---|---|---|---|---|---|---|---|---|---|
| *Chr05* | *19.0* | *19.1* | *6.85E-03* | *-0.8579* | *1.73E-02* | *0.8599* | *6.58E-03* | *-0.3061* | *0.0393* | *0.6204* | *0.5518* | *1.1660* |
| *Chr05* | *20.5* | *20.6* | *8.09E-03* | *-0.7820* | *1.51E-02* | *0.6220* | *5.11E-03* | *-0.5763* | *0.3690* | *0.6616* | *0.2057* | *1.1983* |
| *Chr05* | *20.6* | *20.7* | *9.25E-03* | *-1.1051* | *1.89E-02* | *0.6221* | *6.60E-03* | *-0.3773* | *0.2864* | *0.6498* | *0.7279* | *0.9994* |
| *Chr05* | *20.7* | *20.8* | *9.01E-03* | *-0.7366* | *1.70E-02* | *0.6469* | *5.87E-03* | *-0.5080* | *0.3480* | *0.6538* | *0.2286* | *1.1549* |
| *Chr05* | *20.9* | *21* | *1.04E-02* | *-0.9035* | *2.12E-02* | *0.7789* | *7.24E-03* | *-0.6525* | *0.3008* | *0.6584* | *0.2510* | *1.4314* |
| *Chr05* | *21.0* | *21.1* | *9.32E-03* | *-0.5868* | *1.72E-02* | *0.4935* | *7.12E-03* | *-0.2878* | *0.2364* | *0.5871* | *0.2990* | *0.7813* |
| *Chr05* | *21.1* | *21.2* | *1.08E-02* | *-0.9480* | *2.13E-02* | *0.9625* | *6.33E-03* | *-0.6652* | *0.4126* | *0.7026* | *0.2828* | *1.6277* |
| *Chr05* | *21.2* | *21.3* | *9.03E-03* | *-0.7818* | *1.86E-02* | *0.6586* | *6.10E-03* | *-0.6847* | *0.3242* | *0.6721* | *0.0971* | *1.3433* |
| *Chr05* | *21.3* | *21.4* | *8.09E-03* | *-0.8035* | *1.65E-02* | *0.8474* | *6.32E-03* | *-0.5197* | *0.2194* | *0.6163* | *0.2838* | *1.3671* |
| *Chr05* | *21.4* | *21.5* | *8.38E-03* | *-0.8894* | *1.49E-02* | *0.4240* | *5.22E-03* | *-0.7170* | *0.3764* | *0.6490* | *0.1724* | *1.1410* |
| *Chr05* | *21.5* | *21.6* | *8.41E-03* | *-0.9447* | *1.52E-02* | *0.5291* | *5.13E-03* | *-0.7307* | *0.3895* | *0.6633* | *0.2140* | *1.2598* |
| *Chr05* | *21.8* | *21.9* | *8.27E-03* | *-1.1130* | *1.46E-02* | *0.3985* | *5.23E-03* | *-0.6611* | *0.3681* | *0.6417* | *0.4519* | *1.0595* |
| *Chr05* | *21.9* | *22* | *8.32E-03* | *-0.8841* | *1.85E-02* | *0.6626* | *5.52E-03* | *-0.7310* | *0.3360* | *0.7016* | *0.1531* | *1.3936* |
| *Chr05* | *22.0* | *22.1* | *8.83E-03* | *-0.9598* | *1.97E-02* | *0.6896* | *6.28E-03* | *-0.5790* | *0.2881* | *0.6813* | *0.3809* | *1.2686* |
| *Chr05* | *22.1* | *22.2* | *1.16E-02* | *-0.5849* | *1.79E-02* | *0.4953* | *7.19E-03* | *-0.5452* | *0.3828* | *0.5988* | *0.0398* | *1.0405* |
| *Chr05* | *22.2* | *22.3* | *1.01E-02* | *-1.0096* | *1.91E-02* | *0.2610* | *6.04E-03* | *-0.6482* | *0.4031* | *0.6838* | *0.3615* | *0.9091* |
| *Chr05* | *22.4* | *22.5* | *5.89E-03* | *-0.6253* | *1.76E-02* | *0.7104* | *4.28E-03* | *-0.4972* | *0.2731* | *0.7565* | *0.1281* | *1.2076* |
| *Chr05* | *22.5* | *22.6* | *8.51E-03* | *-0.9605* | *1.71E-02* | *0.6110* | *6.00E-03* | *-0.6379* | *0.2952* | *0.6503* | *0.3227* | *1.2489* |
| *Chr05* | *22.7* | *22.8* | *7.69E-03* | *-1.1362* | *1.50E-02* | *0.4191* | *5.37E-03* | *-0.6258* | *0.3021* | *0.6413* | *0.5103* | *1.0449* |
| *Chr05* | *22.8* | *22.9* | *7.56E-03* | *-0.8436* | *2.09E-02* | *0.4939* | *6.60E-03* | *-0.5422* | *0.1267* | *0.6844* | *0.3013* | *1.0362* |
| ***Chr05*** | ***22.9*** | ***23*** | ***2.84E-03*** | ***-0.9593*** | ***1.26E-02*** | ***0.5275*** | ***2.27E-03*** | ***-0.7795*** | ***0.1999*** | ***0.8199*** | ***0.1799*** | ***1.3070*** |
| ***Chr05*** | ***23.0*** | ***23.1*** | ***4.47E-03*** | ***-0.9123*** | ***1.54E-02*** | ***0.9902*** | ***3.67E-03*** | ***-0.7526*** | ***0.1789*** | ***0.7614*** | ***0.1596*** | ***1.7428*** |
| *Chr05* | *23.1* | *23.2* | *7.07E-03* | *-0.7575* | *1.48E-02* | *0.7022* | *5.28E-03* | *-0.5770* | *0.2532* | *0.6424* | *0.1805* | *1.2792* |
| *Chr05* | *23.2* | *23.3* | *9.80E-03* | *-0.5932* | *1.90E-02* | *0.6240* | *5.74E-03* | *-0.6142* | *0.4141* | *0.6979* | *0.0210* | *1.2382* |
| *Chr05* | *23.3* | *23.4* | *7.78E-03* | *-0.9172* | *1.49E-02* | *0.6325* | *5.19E-03* | *-0.6830* | *0.3334* | *0.6524* | *0.2342* | *1.3155* |
| *Chr05* | *23.4* | *23.5* | *8.88E-03* | *-0.9641* | *1.65E-02* | *0.2895* | *5.48E-03* | *-0.7094* | *0.3828* | *0.6674* | *0.2547* | *0.9989* |
| *Chr05* | *23.5* | *23.6* | *9.51E-03* | *-0.8395* | *1.79E-02* | *0.8107* | *5.24E-03* | *-0.5285* | *0.4493* | *0.7081* | *0.3110* | *1.3392* |
| *Chr05* | *23.9* | *24* | *4.81E-03* | *-1.0835* | *1.65E-02* | *0.4813* | *5.06E-03* | *-0.2477* | *-0.0520* | *0.6936* | *0.8358* | *0.7290* |
| *Chr05* | *24.2* | *24.3* | *9.75E-03* | *-0.7647* | *1.69E-02* | *0.5687* | *7.22E-03* | *-0.2108* | *0.2600* | *0.5739* | *0.5539* | *0.7795* |

| | | | | | | | | | | | | |
|---|---|---|---|---|---|---|---|---|---|---|---|---|
| Chr05 | 28.1 | 28.2 | 5.22E-03 | -1.3893 | 1.66E-02 | 0.4331 | 6.49E-03 | -0.3163 | -0.2426 | 0.6097 | 1.0730 | 0.7494 |
| Chr05 | 28.3 | 28.4 | 5.02E-03 | -1.4314 | 1.52E-02 | 0.6841 | 6.15E-03 | -0.4479 | -0.2261 | 0.5949 | 0.9835 | 1.1319 |
| Chr05 | 28.4 | 28.5 | 5.64E-03 | -1.4265 | 1.68E-02 | 0.3441 | 6.06E-03 | -0.2623 | -0.0749 | 0.6391 | 1.1642 | 0.6064 |
| Chr05 | 30.2 | 30.3 | 9.10E-03 | -1.1471 | 1.63E-02 | 0.2585 | 6.66E-03 | -0.3061 | 0.2676 | 0.5904 | 0.8410 | 0.5646 |
| Chr05 | 51.8 | 51.9 | 3.13E-03 | -0.9787 | 1.62E-02 | 0.4520 | 4.62E-03 | -0.2690 | -0.4732 | 0.7146 | 0.7097 | 0.7210 |
| Chr06 | 11.2 | 11.3 | 1.73E-04 | 0.1156 | 1.70E-03 | 1.4178 | 1.50E-04 | -0.4363 | 0.1311 | 0.9115 | 0.5519 | 1.8541 |
| Chr07 | 15.9 | 16 | 2.01E-02 | 0.5286 | 1.83E-02 | 0.5411 | 6.27E-03 | -0.4871 | 0.6883 | 0.6568 | 1.0158 | 1.0282 |
| Chr07 | 16.3 | 16.4 | 1.48E-02 | 0.6169 | 1.47E-02 | 0.4370 | 5.03E-03 | -0.3649 | 0.6605 | 0.6572 | 0.9818 | 0.8019 |
| Chr07 | 16.4 | 16.5 | 1.47E-02 | 0.6080 | 1.55E-02 | 0.6391 | 5.04E-03 | -0.3763 | 0.6576 | 0.6746 | 0.9843 | 1.0154 |
| Chr07 | 16.6 | 16.7 | 1.43E-02 | 0.2580 | 1.66E-02 | 0.5088 | 6.79E-03 | -0.3055 | 0.5265 | 0.5914 | 0.5635 | 0.8143 |
| Chr07 | 16.7 | 16.8 | 1.85E-02 | 0.5833 | 1.68E-02 | 0.6262 | 5.46E-03 | -0.3097 | 0.7041 | 0.6756 | 0.8931 | 0.9359 |
| Chr07 | 18.3 | 18.4 | 1.37E-02 | 0.5869 | 1.63E-02 | 0.5251 | 5.66E-03 | -0.7695 | 0.5871 | 0.6532 | 1.3564 | 1.2946 |
| Chr07 | 19.4 | 19.5 | 1.75E-02 | 0.9354 | 1.95E-02 | 0.5488 | 6.47E-03 | -0.4398 | 0.6307 | 0.6684 | 1.3752 | 0.9886 |
| Chr07 | 19.5 | 19.6 | 1.81E-02 | 0.8649 | 9.69E-03 | 0.4313 | 4.62E-03 | -0.4165 | 0.7441 | 0.5229 | 1.2814 | 0.8478 |
| Chr07 | 20.0 | 20.1 | 1.29E-02 | 0.0823 | 1.69E-02 | 0.5961 | 6.92E-03 | -0.3812 | 0.4649 | 0.5907 | 0.4635 | 0.9772 |
| Chr07 | 20.6 | 20.7 | 1.20E-02 | 0.6948 | 2.04E-02 | 0.4933 | 7.10E-03 | -0.4671 | 0.4094 | 0.6512 | 1.1619 | 0.9604 |
| Chr07 | 21.0 | 21.1 | 1.91E-02 | 0.1509 | 1.79E-02 | 0.4950 | 5.66E-03 | -0.6470 | 0.7042 | 0.6835 | 0.7980 | 1.1421 |
